# Ecogenomic Diversity of *Clavibacter nebraskensis* in North America

**DOI:** 10.64898/2026.02.11.705226

**Authors:** Luis Fernando Flores-López, Dante M. Callejas, Anne K. Vidaver, Devanshi Khokhani, Oscar Morales-Galván, Verónica Román-Reyna

## Abstract

Goss’s wilt and leaf blight of maize is caused by *Clavibacter nebraskensis* and has reemerged as an important disease in North America. Despite its epidemiological relevance, this species remains poorly characterized in terms of population structure, functional diversity, and ecological differentiation, particularly among strains reported from Mexico. In this study, long-read whole-genome sequencing and phenotypic assays were used to characterize genomic diversity, virulence, and fitness-associated traits in *C. nebraskensis*. We generated 24 long-read genomes, including 20 contemporary Mexican isolates and four historical United States strains collected between 1969 and 1996, and compared them with publicly available genomes from North America and South Africa. Phylogenomic analyses confirmed that all strains cluster within the *C. nebraskensis* clade, and gene accumulation curves supported a closed pangenome with accessory gene variation linked to geographic origin and isolation period. Functional assays showed strain-level variation in virulence, enzymatic activity, bacteriocin antagonism, polysaccharide production, biofilm formation, and pigmentation. Cellulolytic activity was associated with disease severity, whereas pigment-related traits were linked to thiamine metabolism. Overall, these results indicate that *C. nebraskensis* comprises ecologically heterogeneous populations, structured around alternative survival and competition strategies. Integrating genome-wide comparisons with functional characterization of fitness-related traits provides a framework for understanding the biological factors underlying Goss’s wilt. dynamics

## INTRODUCTION

*Clavibacter nebraskensis* (formerly *Corynebacterium nebraskense*; *Clavibacter michiganensis* subsp. *nebraskensis*) is the causal agent of Goss’s wilt and leaf blight of maize, one of the most damaging bacterial diseases of this crop in North America (Jackson et al. 2007; Jardine and Claflin 2016). It was first detected in Dawson County, Nebraska, in 1969, and rapidly spread across the Central Plains during the early 1970s, where it caused severe epidemics before its apparent decline by the mid-1990s (Schuster 1972; Shepherd et al. 2017; Vidaver et al. 1981; Wysong et al. 1973, 1981). The disease reemerged in 2004 in the U.S. Corn Belt, concurring with shifts toward no-till and continuous maize cropping systems (Jackson et al. 2007; Shepherd et al. 2017). Since then, *C. nebraskensis* has expanded its distribution across the United States and Canada and has more recently been reported in Mexico and South Africa, highlighting its increasing global importance (Agarkova et al. 2011; Coertze et al. 2025; Flores-López et al. 2024, 2025; Friskop et al. 2014; Harding et al. 2018; Hosack et al. 2016; Howard et al. 2015; Korus et al. 2011; Malvick et al. 2010; Ruhl et al. 2009; Tambong et al. 2015).

Despite its epidemiological relevance and re-emergence as a major constraint to maize production, *C. nebraskensis* remains one of the least characterized Gram-positive phytopathogens. Foundational aspects of its biology, including virulence mechanisms, secretion systems, metabolic plasticity, population structure, and ecological interactions, remain poorly understood, especially when compared with *C. michiganensis* (*Cm*). *Cm* is the best characterized *Clavibacter* species regarding pathogenicity islands, plasmids that encode effectors, and secreted CAZyme repertoires (Hernández-Aranda et al. 2025; Park et al. 2025; Peritore-Galve et al. 2021; Peritore-Galve et al. 2019; Tancos et al. 2018; Thapa et al. 2017, 2019).

Prior to 2024, only 13 *C. nebraskensis* genomes were available in the National Center for Biotechnology Information (NCBI) (NCBI 2025). Recently, this number increased to 52 genomes, driven by research efforts following the re-emergence and geographic expansion of Goss’s wilt. This expansion includes 46 high-quality long-read assemblies from the United States, Mexico, and South Africa, as well as two short-read genomes from Canada and one genome lacking metadata. Previous comparative genomic studies of *C. nebraskensis* have focused on strains from the United States and Canada, leaving the genetic diversity of Mexican populations underrepresented (Park et al. 2025; Tambong et al. 2015, 2016; Veregge et al. 2025). Two major gaps remain: genomic resources are scarce for historical strains from the early outbreaks (1969–1996), including the first isolates from 1969, and available Mexican genomes have not been fully integrated into comparative genomic or phenotypic analyses. As a result, the evolutionary placement, population structure, and accessory gene diversity of Mexican strains remain poorly resolved, limiting broader inferences about the species across its geographic range.

In Gram-positive phytopathogenic bacteria, virulence and ecological fitness are strongly influenced by secreted molecules, including carbohydrate-active enzymes (CAZymes), extracellular proteases, and bacteriocins. In *Clavibacter* species, secretion through the Sec and Tat pathways mediates host cell wall degradation, nutrient acquisition, and microbial competition. Secreted CAZymes such as cellulases and pectate lyases are well-established contributors to symptom development in related species, particularly for *C. michiganensis* (Hernández-Aranda et al. 2025; Park et al. 2025; Sen et al. 2015; Stevens et al. 2021; Tancos et al. 2018; Thapa et al. 2017). Despite this, prior studies have not systematically examined whether variation in these traits, including CAZymes, bacteriocins, extracellular enzymes, and other social behaviors, correlates with differences in *C. nebraskensis* virulence. Bacteriocins have historically been described only through bacterial growth inhibition assays and strain-typing schemes, without identification of their genetic basis or evaluation of their ecological roles during plant colonization (Gross and Vidaver 1979; Smidt and Vidaver 1986; Vidaver et al. 1981).

Here, we address these gaps by generating genomic resources for *C. nebraskensis* through long-read sequencing of 24 strains, including 20 contemporary Mexican isolates and four historical U.S. strains spanning 1969–1996, and by comparing them with 15 publicly available genomes from North America and South Africa. We then focused on functional analyses of the 24 sequenced strains to integrate high-quality genomic data with standardized phenotypic assays, including virulence on maize, enzymatic activity profiling, bacteriocin-mediated antagonism, pigment production, and growth kinetics in maize apoplastic fluid. By linking genomic variation to functional phenotypes, this study provides the first ecological–evolutionary framework connecting population structure, secreted molecules, and virulence diversity in *C. nebraskensis*.

## MATERIALS AND METHODS

### Bacterial strains and growth conditions

A total of 25 *Clavibacter nebraskensis* strains were included in this study (Table 1). These comprised 24 strains sequenced in this study and the type strain NCPPB 2581. The 24 strains include, 20 contemporary Mexican isolates obtained from symptomatic maize plants in Panotla, Tlaxcala, and Saltillo, Coahuila (Flores-López et al. 2024) and four historical U.S. isolates collected between 1969 and the late 1990s. The type strain NCPPB 2581, also refereed as FH36, was included as a phenotypic reference but was not re-sequenced. All 25 strains were routinely cultured on Nutrient Broth-Yeast Extract (NBY) medium at 25 °C for 48–72 h prior to downstream analyses (Vidaver 1967).

### Pathogenicity assays on maize plants

All *C. nebraskensis* strains were grown overnight in liquid NBY broth at 25 °C with shaking at 180 rpm. Then the cultures were transferred to fresh broth cultures and grown to mid-log phase. Cells were harvested by centrifugation at 8,000 g, washed once with sterile distilled water, and resuspended in phosphate-buffered saline (PBS) (Shepherd et al. 2017). Bacterial suspensions were adjusted to 10^8^ CFU mL^-1^, corresponding to OD_640_= ∼0.7. PBS was used as the mock inoculation control, and the type strain NCPPB2581 was included as a pathogenicity reference.

Virulence was assessed using the maize inbred line A632, grown in a controlled-environment growth chamber (Percival Scientific) under a 16 h light and 8 h dark photoperiod, 25 °C, and 85% relative humidity. Plants were inoculated at the V4 developmental stage. A fully expanded leaf on each plant was infiltrated at mid-leaf using a needleless syringe. Immediately after infiltration, the infiltrated area was marked with a permanent ink marker to enable precise quantification of lesion expansion. Each strain was inoculated in three leaves. To enhance early disease development, plants were maintained for the first 72 h post-inoculation under continuous mist generated by an ultrasonic humidifier within the growth chamber (SPT SU-4010). Thereafter, growth chamber remained at 85% RH throughout the assay. Lesion expansion was measured at 4, 8, and 12 days post-inoculation (dpi) in both proximal and distal directions. Leaf length was measured at 12 dpi to normalize lesion development across plants.

The lesion percentage was calculated as:

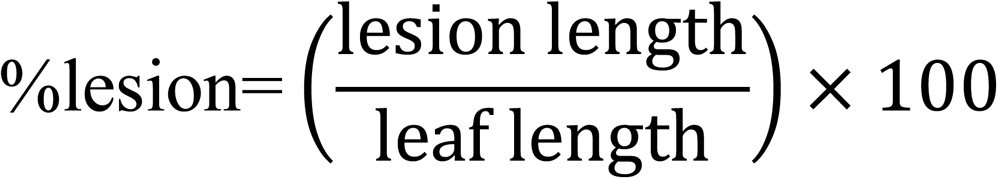

Lesion measurements were processed in R v4.3 (R Core Team 2024). Percentage lesion values were used to compute disease progress curves and the area under the disease progress curve (AUDPC) using the trapezoidal method (DescTools::AUC). Differences among strains were evaluated using one-way ANOVA, followed by Tukey’s HSD for multiple comparisons. Data visualization included disease progress curves (mean ± SD) and AUDPC boxplots annotated with significance group letters using ggplot2 (Wickham 2016).

### Genomic DNA extraction and quality control

High-molecular-weight genomic DNA was extracted from each *C. nebraskensis* strain pure culture using the Wizard® HMW DNA Extraction Kit (Promega, Cat. A2920), following the manufacturer’s protocol for Gram-positive bacteria with slight modifications. Briefly, overnight broth cultures were grown at 25 °C with shaking (180 rpm), cells were harvested by centrifugation at 8,000 g, and resuspended in a lysozyme solution (10 mg/mL) and incubated for 1 hour to weaken the cell wall. Lysis and DNA precipitation were performed following the manufacturer’s protocol. DNA was rehydrated overnight to preserve long-fragment integrity. DNA concentration was measured using a Quantus Fluorometer (Promega) with the QuantiFluor dsDNA System. Purity ratios (A260/280 and A260/230) were examined spectrophotometrically in a UV5Nano (Metler Toledo, Columbus, OH), and DNA integrity (high-molecular-weight fragments) was assessed on 0.8% agarose gels.

### Library preparation and sequencing

Sequencing libraries were prepared using the Oxford Nanopore Technologies (ONT) Native Barcoding Kit 24 V14 (SQK-NBD114.24) according to the manufacturer’s ligation-based native barcoding protocol. For each strain, 400 ng of HMW DNA was subjected to end-repair and dA-tailing, followed by ligation of individual native barcodes. Barcoded samples were pooled equimolarly into a single multiplexed library and finalized with adaptor ligation using reagents provided with the V14 chemistry. All library-prep steps were performed using low-bind consumables to maintain long-read performance. Multiplexed libraries were loaded onto MinION R10.4.1 flow cells (FLO-MIN114) and sequenced on a MinION Mk1B device using MinKNOW default settings for ligation-based libraries. Runs were conducted for 48 hours or until pore activity dropped below recommended thresholds. Basecalling and demultiplexing were performed using Dorado v1.0.0 in high-accuracy mode, with an HAC model, generating FASTQ files for downstream analysis.

### Genome assembly, rotation, completeness, and Annotation

Demultiplexed Nanopore reads were assembled with Flye v2.9.5 using the parameters nano-hq, genome size 3g, three iterations, and a read error of 0.03 (Kolmogorov et al. 2019). All genomes were rotated to position the *dnaA* gene at the origin of replication. Rotation was performed using a Python script provided by Dr. Ralf Koebnik (IRD, France), which identifies the *dnaA* locus and reorders the sequence so that *dnaA* marks the first base of the genome. Genome completeness was evaluated using BUSCO v6.0.0 with the actinomycetota_odb12 lineage dataset (Tegenfeldt et al. 2025). BUSCO was run to quantify the proportion of single-copy orthologs classified as complete, duplicated, fragmented, or missing. Finally, genome annotation was performed using Bakta Web with default parameters (Beyvers et al. 2025). Annotated genomes were exported in FASTA, GFF3, and GenBank formats for downstream comparative genomics, pangenome analysis, and gene mining.

### Phylogenomic reconstruction

Phylogenomic relationships were reconstructed using PHANTASM, an alignment-free, whole-genome distance approach based on k-mer frequencies (Wirth and Bush 2023). The dataset included all 39 *C. nebraskensis* genomes (Table S1) to place the Mexican and U.S. isolates within the full taxonomic breadth of the genus. PHANTASM was run with default parameters to compute pairwise genomic distances, and a neighbor-joining tree was generated from the resulting matrix. The phylogeny was exported to iTOL v6, where it was visually refined and annotated with metadata layers including geographic origin.

### Pangenome analysis

The pangenome was reconstructed using Panaroo v1.3, a graph-based pipeline optimized for long-read assemblies and frameshift correction (Tonkin-Hill et al. 2020). Annotated GFF3 files generated by Bakta served as inputs, and Panaroo was executed with the clean-mode strict to minimize spurious gene calls. The resulting gene presence-absence matrix was imported into R for downstream analysis, in which gene clusters were converted into a binary matrix to classify core, shell, and cloud components. Pangenome openness was evaluated using Heaps’ Law, implemented through 100 randomized genome permutations and non-linear fitting (nlsLM) to estimate the γ parameter. Gene intersection patterns across all genomes were visualized using UpSetR, and accumulation curves were plotted to quantify novel gene contributions and genome-wide diversity (Conway et al. 2017).

### Maize Apoplast Growth Assays

Bacterial growth kinetics in maize apoplast were evaluated using a 96 microplate-based assay. Apoplastic fluid was extracted from healthy maize leaves by vacuum infiltration with sterile distilled water followed by centrifugal recovery (Roman-Reyna and Rathjen 2017). The recovered fluid was diluted 1:1 (v/v) in sterile distilled water and passed through a 0.22 µm polyethersulfone (PES) membrane filter to eliminate microbial contaminants.

Each strain was cultured overnight in NBY at 25 °C with agitation (120 rpm). Cultures were pelleted, washed once in sterile distilled water, and resuspended in PBS. Suspensions were adjusted to an optical density of 0.1 at 640 nm, and 5 μL of each inoculum was dispensed into wells containing 95 μL of sterile maize apoplast solution. Each strain had three technical replicates. Wells only containing apoplast served as blanks. Plates were sealed with a gas-permeable breathable membrane to prevent condensation while maintaining aeration. Growth kinetics were recorded using a SpectraMax i3 multi-mode plate reader configured in kinetic mode at OD_640_. Plates were incubated at 25 °C for 72 hours, with readings performed automatically every hour. Prior to each reading, the plate was subjected to orbital shaking for 30 seconds to ensure homogenization and minimize micro-settling artifacts. The plate experiment was repeated twice. Raw OD₆₄₀ time-series data were imported into R for processing using gcplyr (Blazanin 2024). After reshaping into tidy format and mapping wells to strain identities, background correction was applied using either the time-resolved mean OD of blank wells or, when absent, an early-timepoint per-well baseline. Curves were smoothed using a moving median filter, followed by loess regression, to generate high-resolution growth trajectories. Per-capita growth rates (µ, h⁻¹) were derived from the first derivative of log-transformed smoothed OD values. Diauxic shifts were identified for each well by locating the first local maximum in µ followed by a local minimum, defining the diauxic break (t_break_). Growth parameters were extracted separately for the first (P1; inoculation to t_break_) and second (P2; t_break_ to 72 h) phases. Metrics included the maximum per-capita growth rate (µ_max_), doubling time (dt, in minutes), carrying capacity (K), initial OD (N₀), and area under the curve (AUC). For each strain, median values from the six replicates were used for downstream comparisons. Statistical analyses were performed by rank-based nonparametric methods. Differences among strains in doubling time, AUC, K, and t_break were evaluated separately for each phase using Kruskal–Wallis tests, followed by Dunn’s post hoc tests with Benjamini–Hochberg correction. Compact letter displays were generated using multcompView, and figures were constructed using ggplot2.

### Enzymatic activity and genetic comparisons for cellulose-degrading ability

#### Cellulase activity in vitro

The cellulose-degrading ability of *C. nebraskensis* strains was evaluated using a carboxymethyl cellulose (CMC) plate assay, adapted from Pfeilmeier et al., (2024). Cellulase-inducing medium (RSD–CMC) was prepared according to the formulation described by Davis and Vidaver (2001) as described in the Laboratory Guide for Identification of Plant Pathogenic Bacteria (Schaad et al., 2001). The basal RSD agar per liter included: 1.0 g yeast extract, 1.0 g (NH₄)₂HPO₄, 1.0 g MgSO₄·7H₂O, 0.2 g KCl, 1.0 g L-cysteine, 15 mL bovine hemin chloride (0.1% w/v in 0.5 N NaOH), 10 mL BSA fraction V (20% sterile solution), and 15 g agar. Sodium CMC (0.5% w/v; Sigma-Aldrich C5678) was added as the only carbon source prior to autoclaving. The inoculum was prepared by culturing each strain overnight in NBY at 25 °C with shaking at 120 rpm. The cultures were then centrifuged at approximately 1,000 × g for 10 minutes, washed once, and resuspended in PBS to an optical density of 0.1 at 640 nm. Plates were spotted in triplicate with 7 µL of these standardized suspensions and incubated at 25 °C for 10 days.

After incubation, cellulose degradation was detected by flooding the plates with 0.1% (w/v) Congo red for 30 minutes and then washing repeatedly with 1 M NaCl until the background was clear and hydrolysis zones appeared. The presence of yellow halos around colonies indicated carboxymethyl cellulase activity. Halo diameters (mm) were measured from digital images of the plates with ImageJ, and their areas (cm²) were calculated using π×(d/2)² (Schneider et al. 2012). All analyses were conducted using R v4.4.2. For each strain, halo areas served as response variables measured across technical replicates. Normality and homogeneity of variances were assessed visually, followed by a one-way ANOVA with strain as the fixed factor. Significant differences were further explored with Tukey’s HSD test (α = 0.05). Results were visualized through boxplots and compact letter displays that highlight differences among strains, created with ggplot2 (Wickham, 2016).

#### Cellulase-associated gene comparisons

To investigate the genetic basis underlying variation in cellulase activity, the focus was on two genes involved in cellulose degradation. One was *exlX1*, a gene previously associated with cellulase activity and virulence in *C. nebraskensis* (Webster et al., 2024). The second gene, a predicted secreted endoglucanase (glycoside hydrolase family) present in all genomes and showing sequence polymorphism, was identified. For each gene, DNA sequences were extracted from all genomes and aligned using MUSCLE in MEGA v12 under default parameters. Alignments were visually inspected to verify preservation of the reading frame and trimmed to shared coding regions across all strains. The two trimmed alignments were concatenated with the software Concatenator and exported in NEXUS format for haplotype reconstruction (Vences et al. 2022). PopART was employed to generate a haplotype network using the median-joining algorithm (Leigh and Bryant 2015). Additionally, haplotype reconstruction, nucleotide diversity (π), haplotype diversity (Hd), and Tajima’s D were calculated. The resulting haplotype network was visualized by coloring haplotypes according to country of origin.

### Enzymatic activity and genetic comparisons for starch

#### Amylase activity *in vitro*

Starch hydrolysis was evaluated using soluble starch agar, following a method previously used for plant-pathogenic bacteria, with slight modifications (Schaad et al. 2001). Plates were prepared by dissolving 10 g of soluble starch and 16 g of agar in 1 L of distilled water, then autoclaved and poured into Petri dishes. Each *C. nebraskensis* strain was grown overnight in NBY broth, adjusted to an OD₆₄₀ of 0.1 in PBS, and spot-inoculated with 7 µL onto the starch agar plates in triplicate. The plates were incubated at 25 °C for 10 days. Starch degradation was detected by flooding the plates with Lugol’s iodine solution, which stains intact starch dark blue-black. Amylase activity appeared as clear yellow halos around the colonies.

#### Identification of starch-degrading enzyme genes

To identify genes potentially responsible for starch hydrolysis in *C. nebraskensis*, genome mining was performed using biochemical models for bacterial starch degradation (Hii et al. 2012; Lin et al. 2021a; Mehta and Satyanarayana 2016). A list of genes coding for amylolytic enzymes such as α-amylase (EC 3.2.1.1), pullulanase (EC 3.2.1.41), neopullulanase (EC 3.2.1.135), and α-glucosidase (EC 3.2.1.20), was defined based on the enzymatic roles in α-glucan metabolism and functional annotations from CAZy and BRENDA. Bakta-annotated files were used to extract the sequences associated with the EC numbers or having the enzyme names. This process resulted in a curated set of starch-degrading enzyme annotations for each genome, which were saved as strain-specific summary tables. To compare starch-degradation gene content across strains, copy numbers were summarized for each annotation description across all genomes. Data processing and visualization were performed in R (v4.4.2). Annotation descriptions were grouped into four functional categories based on keyword-based classification of Bakta product descriptions: endo-α-amylase (GH13-like), GlgX-like debranching enzymes, neopullulanase-like dual-specificity enzymes, and glucoamylase-like exo-amylases. Copy number per strain and annotation was visualized as a dot-matrix, with point size proportional to gene copy number and color indicating functional group, and strains ordered according to virulence ranking.

### Enzymatic activity and genetic comparisons for proteases

#### Protease hydrolysis assay

Protease activity was evaluated using a nutrient gelatin hydrolysis medium formulated (per liter) with peptone 2 g, beef extract 1 g, NaCl 5 g, gelatin 6 g, and agar 12 g. Each strain was grown overnight in NBY broth at 25 °C, pelleted, washed once with sterile distilled water, and resuspended in phosphate-buffered saline (PBS) to an OD₆₄₀ of 0.1. Each strain was spotted in 7-µL drops, in triplicate, and incubated at 25 °C for 10 days. After incubation, proteolytic activity was visualized by flooding plates with acidic mercuric chloride reagent (per 100 mL: 12 g mercuric chloride dissolved in 80 mL distilled water, then supplemented with 16 mL concentrated HCl) (Klement et al. 1990). This reagent precipitates undegraded gelatin, producing an opaque background. Plates were flooded for several minutes, rinsed thoroughly with distilled water, and allowed to air-dry. To stabilize and enhance contrast of the hydrolysis halos, plates were then overlaid with 1–2 mL of Loeffler’s methylene blue for 5 min and rinsed with 70% ethanol, which fixed the stained matrix and accentuated clear zones around colonies. Protease activity was quantified as the diameter of each halo. Halo areas were calculated and analyzed in R v4.4.2 using a one-way ANOVA with strain as the fixed factor. Assumptions of normality and homogeneity of variances were verified graphically. When significant effects were detected (α = 0.05), a Tukey’s HSD was applied to resolve pairwise differences. Compact letter displays were generated using the multcompView package in R (Piepho 2004).

#### Comparative analysis of protease-coding genes

To investigate the genetic diversity of proteolytic enzymes that may contribute to gelatin hydrolysis in *C. nebraskensis*, a comparative analysis of 22 protease-coding genes conserved across all genomes was conducted. The selection of genes was guided by curated enzyme classifications in the MEROPS peptidase database and entries from BRENDA/Expasy, with a focus on proteases in the EC 3.4 class, which comprises functionally characterized serine proteases, metalloproteases, and other endopeptidases involved in extracellular protein degradation (Feuilloley et al. 2024, 2024; Rawlings et al. 2018; Toth and Fridman 2001). Candidate loci were identified from Bakta annotations by querying product descriptions and EC numbers associated with proteolytic families, including subtilisin-like serine proteases (MEROPS family S8), thermolysin-like metallopeptidases (M4), and serralysin-related metalloproteases (M10). To ensure consistency in analysis across genomes, only those proteases genes found in all strains were retained for downstream comparative analyses.

Protease annotations were then categorized into functional classes using a rule-based keyword classifier applied to the Bakta description text. Categories were designed to capture major mechanistic groups relevant to extracellular protein turnover and proteolysis, including virulence-associated trypsin-like serine proteases (Pat-1/Chp/Ppa-like), subtilisin-like serine proteases (S8), thermolysin/gelatinase-like metalloproteases (M4), other metalloproteases, amino- and dipeptidases involved in peptide processing, cysteine proteases, aspartic proteases, ATP-dependent housekeeping proteases (e.g., Clp/Lon/FtsH), and signal peptidases. To compare gene abundance across strains, copy number was summarized for each annotation description in each genome (i.e., the number of occurrences of the same description within a strain). Data processing and visualization were performed in R (version 4.4.2) using the ggplot2 package. Copy number distributions were visualized as a dot matrix, where point size represents copy number and point color indicates protease functional class; strains were ordered by virulence ranking.

### Social behavior

#### Biofilm formation Assay

Biofilm formation was quantified using a static crystal violet assay adapted from Merritt et al., (2006). To assess biofilm formation under host-relevant conditions, assays were performed in maize apoplastic fluid rather than rich laboratory media, consistent with the apoplast-based growth assays described above. Each *C. nebraskensis* strain was cultured overnight in NBY broth at 25 °C with shaking (120 rpm), pelleted, and washed once with sterile distilled water. Cells were resuspended in PBS and standardized to an optical density of OD₆₄₀ = 0.1. For each strain, 5 μL of the standardized inoculum was added to 95 μL of sterile maize apoplastic fluid in flat-bottom 96-well polystyrene microplates (Costar®, Corning Inc., Corning, NY, USA). Apoplastic fluid was extracted from healthy maize leaves via vacuum infiltration and centrifugation, as described in the growth methods.

Plates were incubated at 25 °C for 48 hours to allow biofilm development. After incubation, wells were gently washed three times with sterile distilled water to remove non-adherent cells. Adherent biofilms were stained with 0.1% (w/v) crystal violet for 15 minutes at room temperature, then rinsed three times to remove excess dye. Stained biofilms were solubilized by adding 200 μL of 30% acetic acid per well, and absorbance was measured at 590 nm (OD₅₉₀) using a SpectraMax i3 microplate reader. Biofilm assays were conducted with three technical replicates per strain. Blank-corrected OD values were analyzed in R v4.4.2 using the Kruskal–Wallis test to evaluate overall differences in biofilm formation. Pairwise comparisons among strains were performed using Dunn’s test with Benjamini–Hochberg correction for multiple testing. Data were visualized using ggplot2, with strain-level means, standard deviations, and replicate distributions displayed. Significant groupings were generated using compact-letter displays.

#### Extracellular Polysaccharide (EPS) Production Assays

EPS production was assessed using qualitative and quantitative approaches. All strains were initially cultured on solid NBY medium, then grown overnight in liquid NBY at 25 °C with shaking (300 rpm). Cells were harvested by centrifugation, washed once with phosphate-buffered saline (PBS), and resuspended to an optical density of 0.1 at 640 nm in PBS. Visual detection of EPS was performed on a solid medium supplemented with ruthenium red, following the method described by Yadav et al., (2024) for *Lactobacillus* spp., with modifications. This protocol was selected because it is a validated EPS assay for Gram-positive bacteria, and comparable methods have not been standardized for Gram-positive plant pathogenic bacteria. The basal medium consisted of a potato infusion prepared by boiling 200 g of peeled and chopped potatoes in 1 L of distilled water, filtering the extract, and adjusting the volume back to 1 L. The infusion was supplemented with 20 g/L sucrose and 15 g/L agar, then autoclaved. Once cooled to ∼50 °C, 10 mL/L of sterile-filtered 0.8% (w/v) ruthenium red solution was added. Bacterial suspensions were spotted in triplicate (7 µL per replicate) and incubated at 25 °C for 4 days. Potato infusion was used to minimize pigmentation bias, as *C. nebraskensis* produces orange pigmentation in the presence of thiamine. Unlike potato dextrose agar (PDA), potato infusion allows control of the carbon source and concentration while suppressing pigment production, thereby improving EPS visualization (Bradbury, 1991).

EPS concentration was determined using the phenol–sulfuric acid assay (DuBois et al. 1956). Two liquid culture conditions were tested: (i) potato–sucrose medium as described above, without agar or dye, and (ii) maize apoplast extract prepared as previously described. For each condition, cultures were established using a 95:5 volumetric ratio of sterile medium to bacterial suspension. Specifically, 1,425 µL of medium and 75 µL of inoculum were used for the sucrose medium, while apoplast cultures used 95 µL of apoplast fluid and 5 µL of bacterial suspension. Cultures were set up in 24-well and 96-well plates respectively (Costar®, Corning Inc.), sealed with gas-permeable membranes, and incubated at 25 °C with shaking (300 rpm) for 48 hours. EPS was recovered from culture supernatants by ethanol precipitation. After incubation, cultures were centrifuged at 10,000 × g for 15 minutes at 4 °C. Supernatants were transferred to clean tubes, mixed with three volumes of cold absolute ethanol (e.g., 1 mL supernatant + 3 mL ethanol), and incubated overnight at −20 °C. EPS pellets were collected by centrifugation (10,000 × g, 20 min, 4 °C) and resuspended in distilled water.

To quantify total carbohydrate content, 500 µL of each EPS preparation was mixed with 5% (v/v) phenol and concentrated sulfuric acid following the method of Dubois et al. Absorbance was measured at 490 nm using a SpectraMax i3 microplate reader (Molecular Devices). A glucose standard curve (0–100 µg) was prepared from a 1 mg/mL stock solution and used to estimate EPS concentrations. Quantitative data from the phenol–sulfuric acid assay were normalized to cell density by dividing glucose-equivalent concentrations (µg) by the corresponding optical density of each culture. For each medium (sucrose and apoplast), EPS production normalized per OD was compared among strains using non-parametric statistics. Kruskal–Wallis tests were conducted in R v4.4.2 to assess overall differences among strains. When significant effects were detected (α = 0.05), strain-level comparisons were resolved with Duncan’s multiple range test using the agricolae package. Compact letter displays were generated to denote homogeneous groups. Figures were created with ggplot2 using consistent palettes and strain ordering across all phenotypic assays.

#### Bioinformatic Analysis of Social Trait Genes

Prediction of genes associated with bacterial social behaviors was performed using SOCfinder v1.0.0 (Belcher et al. 2023), a comparative genomics framework designed to identify loci linked to cooperative and collective functions in bacteria. Analyses were conducted using the parameter set optimized for Gram-positive genomes.

SOCfinder output consisted of strain-specific lists of predicted social genes annotated by protein identifiers and KO-based functional descriptions. SOCfinder results were manually curated in Microsoft Excel to standardize protein annotations and verify functional assignments based on KO definitions and associated descriptions. This curation step ensured consistency across strains and removed ambiguous or redundant entries prior to downstream quantitative analyses. Curated SOCfinder data were subsequently analyzed in R (v4.4.2). Social gene investment was quantified for each strain as the proportion of predicted social genes relative to the total number of protein-coding sequences, as determined from Bakta annotations. These values were visualized in a lollipop plot, with strains ordered by virulence rank. In parallel, predicted social genes were classified into functional categories representing distinct social strategies (competition, biofilm/extracellular polysaccharide production, regulation and signaling, protection and stress response, secretion and transport, and unclassified functions) using keyword-based parsing of KO definitions. The distribution of these strategy categories across strains was summarized as gene counts and visualized as a dot-matrix, with point size proportional to gene count and color indicating functional category.

### Bacteriocin

#### Bacteriocin Activity Assay

Bacteriocin production was assessed using a modified overlay method adapted from protocols established for Gram-positive plant pathogenic bacteria (Gross and Vidaver 1979; Nelson and Semeniuk 1964; Vidaver et al. 1981). Two types of agar media were used to evaluate activity: nutrient broth yeast extract (NBY) and Modified Burkholder’s Agar Lacking peptone (MBAL).

Producer strains were cultured overnight in liquid NBY at 25 °C with shaking, standardized to an optical density of 0.1 at 640 nm, and spotted in 7 µL aliquots onto NBY or MBAL agar plates. Each strain was tested in a triplicate. After a 48-hour incubation at 25 °C, colonies were replicated onto fresh plates using a sterile velvet replicator and incubated at 20 °C for an additional four days to allow bacteriocin accumulation. Producers were then inactivated by exposure to chloroform vapors for two hours under a chemical fume hood, followed by evaporation of residual chloroform prior to overlay. Indicator strains were grown overnight, diluted to approximately 10⁸ CFU mL⁻¹, and 100 µL of the suspension was mixed with 2.5 mL of molten soft agar (0.7%) prepared in NBY or MBAL, previously cooled to 45 °C. The mixture was poured over the inactivated producer plates and incubated at 25 °C for 24–48 hours. Bacteriocin activity was recorded as the presence of growth inhibition zones surrounding producer colonies. Inhibition diameters were measured in centimeters, and data were analyzed to assess strain-level differences.

#### Bioinformatic Prediction of Bacteriocins

To identify putative bacteriocin biosynthetic gene clusters (BGCs), the genomes of all strains were analyzed with BAGEL4 using default parameters optimized for Gram-positive bacteria (van Heel et al. 2018). For each strain, both nucleotide and GenBank-formatted outputs were retrieved. Predicted BGCs were examined using clinker to assess amino acid content and synteny at the BGC level across strains (Gilchrist and Chooi 2021). The nucleotide sequences of the predicted bacteriocin genes were concatenated by strain and aligned to reconstruct a phylogenetic tree, allowing visualization of strain-level clustering based on bacteriocin content. These groupings were subsequently compared to phenotypic differences in bacteriocin production observed in the overlay assays to evaluate potential correlations between genomic content and inhibitory activity.

### Pigment-associated environmental protection traits

#### Pigment phenotyping and spectral absorbance

Pigment-associated traits linked to environmental protection were assessed both visually and by absorbance spectroscopy. Overnight liquid cultures were serially diluted and plated on NBY agar to document colony pigmentation. For spectral profiling, cells from liquid cultures were harvested, washed with sterile distilled water, and resuspended in PBS. Cell suspensions were normalized to approximately 1 × 10⁸ cells mL⁻¹ to ensure equivalent biomass across strains and minimize cell-density–driven bias in absorbance measurements. Suspensions were dispensed into 96-well plates and then analyzed by scanning the absorbance in the wavelength region of 200–1000 nm using a SpectraMax i3 microplate reader. Blank wells containing only PBS were included and subtracted wavelength by wavelength. Spectral data were processed in R. Absorbance values were averaged across three technical replicates per strain and wavelength, and spectral curves were visualized using ggplot2. Peak detection was performed using the pracma package. For comparative analyses, absorbance matrices (wavelength × strain) were constructed and used for hierarchical clustering based on Euclidean distances with Ward’s minimum-variance method (Ward.D2), implemented in base stats.

#### Terpene biosynthetic gene clusters

Putative pigment-associated terpene biosynthetic gene clusters were identified using antiSMASH (Blin et al. 2021). Two terpene-associated loci (BGC regions 6 and 9) were detected and extracted as nucleotide sequences. These loci were aligned, trimmed, and concatenated following the same workflow used for other loci in this study. Sequence variation was summarized by reconstructing haplotype networks using the median-joining algorithm in PopART to evaluate genetic diversity across strains.

#### Thiamine-related gene identification and comparative analysis

Genes involved in thiamine biosynthesis, activation, transport, and regulation were identified through genome mining guided by established bacterial thiamine metabolism pathways and functional annotations (Devedjiev et al. 2004; Du et al. 2011; Jurgenson et al. 2009; Kim et al. 2020; Webb et al. 1998). Targeted genes included components of thiamine biosynthesis (e.g., *thiC, thiD, thiE*), thiamine activation and salvage pathways (e.g., *thiM, thiK*), transport systems, and associated regulatory elements. The distribution of thiamine-related genes was summarized as presence/absence and copy-number matrices. Comparative analyses were performed in R to visualize gene content and copy-number variation across strains, facilitating the assessment of lineage-specific patterns potentially linked to pigment production and environmental protection traits.

## Results

### Phylogenomic evidence supports species-level placement of study strains as *C. nebraskensis*

A phylogenomic analysis was conducted to assess the taxonomic placement of the strains analyzed within the genus *Clavibacter*, resolving all described species into well-supported species-level clades consistent with current genomic and taxonomic classifications (Figure 1). All *C. nebraskensis* strains formed a monophyletic group. Within the *C. nebraskensis* clade, strains were distributed across multiple internal branches, indicating genomic diversification within the species.

**Figure 1.**
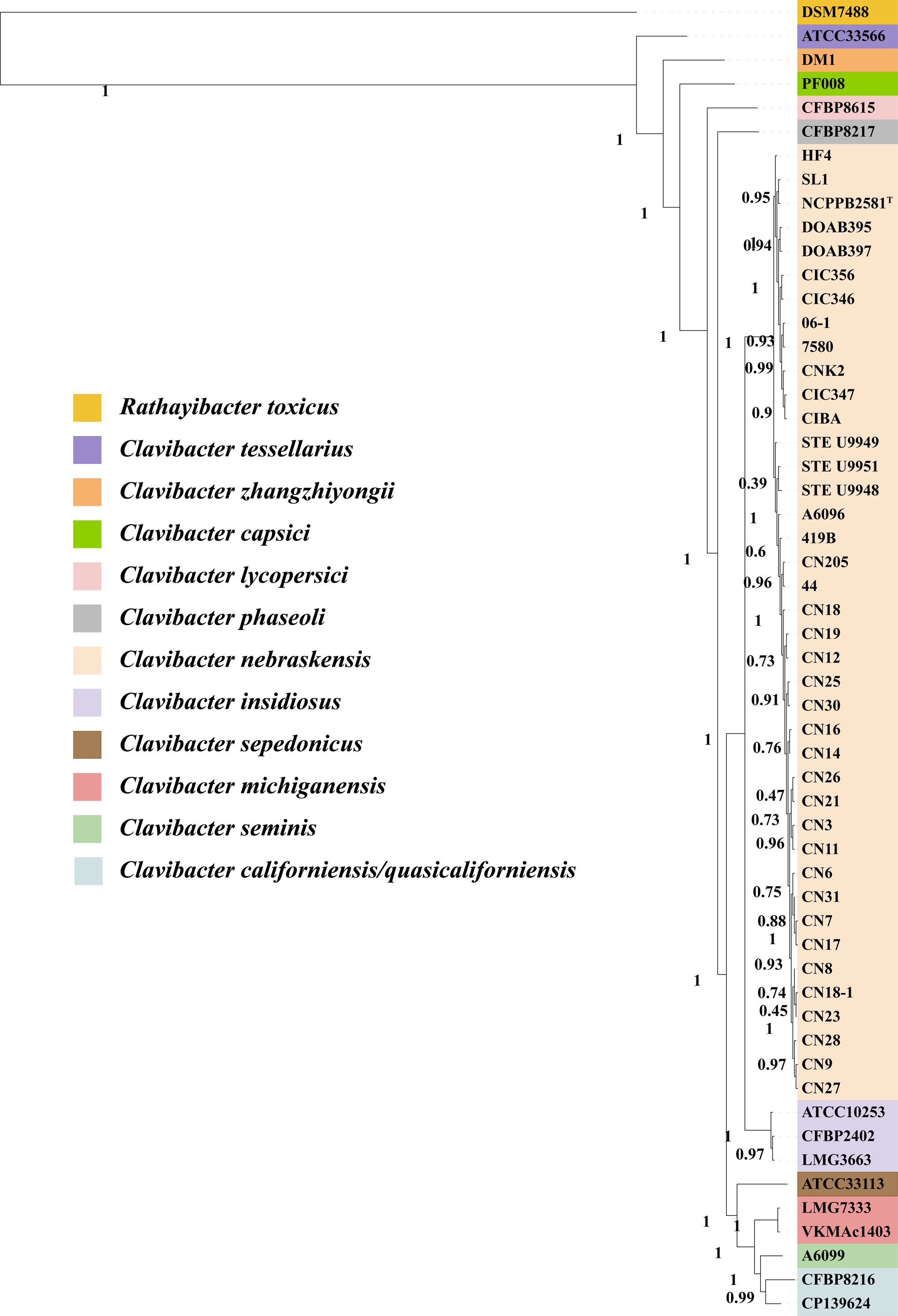
Phylogenomic relationships within the genus *Clavibacter*. Phylogenomic tree inferred with PHANTASM from whole-genome data. Branch colors show species within *Clavibacter*. *Rathayibacter toxicus* DSM 7488 was used as outgroup. Node support values are at internal branches.

### Pangenome structure suggests temporal and geographic patterns within *C. nebraskensis*

*C. nebraskensis* has a close pangenome based on the gene accumulation curve (Supplementary Figure S1). The total number of new genes was steady as more genomes were sequenced, suggesting saturation of the species’ genetic diversity. The *C. nebraskensis* pangenome revealed gene distribution patterns associated with year of isolation and geographic origin (Figure 2). Mexican strains shared more accessory genes with the 1969 U.S. isolate than with the 1971 isolate (39 genes on average). The 1971 isolate instead shared more accessory genes with other U.S. strains (42 genes on average), suggesting that Mexican strains are genetically closer to the 1969 U.S. isolate. Together, these results indicate that variation in the *C. nebraskensis* pangenome reflects differential retention and loss of accessory genes.

**Figure 2.**
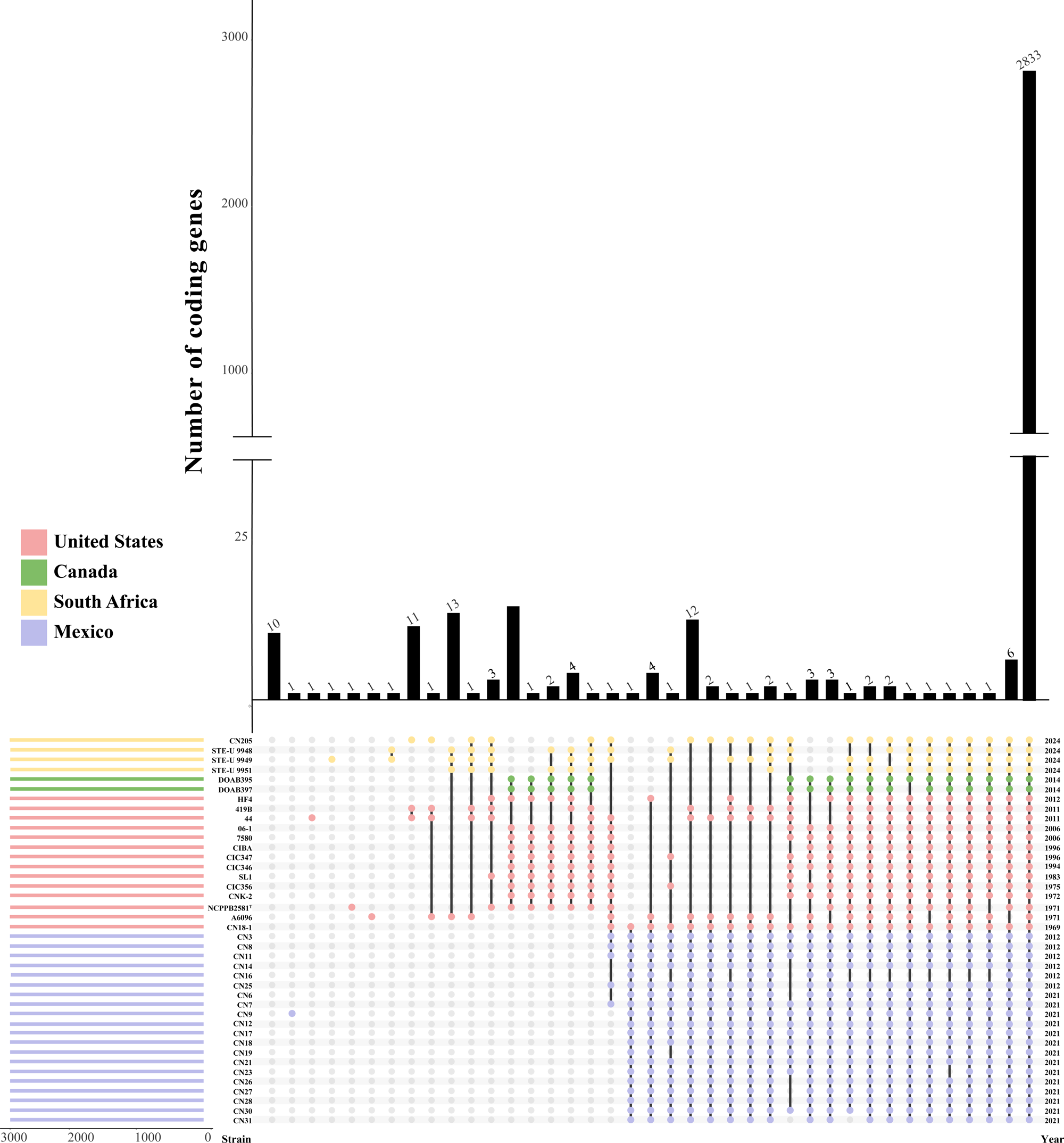
Pangenome structure across *Clavibacter nebraskensis* strains. UpSet plot showing intersections of shared coding genes across strains. Vertical bars represent the number of genes in each intersection, while the connected dots below indicate the specific strain combinations contributing to each set. Intersections correspond to genes shared by all strains (core genome), genes shared by a subset of strains (shell genome), and strain-specific genes (cloud genome). Strains are color-coded by country of origin, as indicated in the legend. The year of isolation for each strain is shown on the right.

These pangenome patterns indicated that the Mexican strains represent a distinct *C. nebraskensis* group. Therefore, to link genomic differentiation to biologically relevant outcomes, phenotypic characterization was prioritized for the Mexican strains while retaining five reference U.S. isolates as historical comparators.

### Disease severity across *C. nebraskensis* strains

The disease progress over time and the area under the disease progress curve (AUDPC) were significantly different among *C. nebraskensis* strains (one-way ANOVA: F = 6.229, P = 1.49 × 10⁻⁸) (Figure 3A) when using the cultivar A632. The AUDPC values varied across strains, with CN21(MX, 2021) exhibiting the highest disease severity (485 %·day), followed by CN18-1 (USA, 1969) (449 %·day) and CN28 (MX, 2021) (442 %·day). Intermediate disease levels were observed for multiple strains, whereas NCPPB2581^T^ (USA, 1971), CIC346(USA, 1994), CIC356 (USA, 1975), and CIC347 (USA, 1996) consistently showed low AUDPC values, with CIC347 displaying the lowest disease severity among inoculated strains (12.9 %·day). In addition to quantitative differences in lesion expansion, strains differed in symptom types (Figure 3B). Observed symptoms include water soaking, discrete foliar freckling, tissue reddening, and extensive necrosis. Some strains caused localized water-soaked or freckled lesions, while others caused reddening of leaf tissue and large necrotic areas.

**Figure 3.**
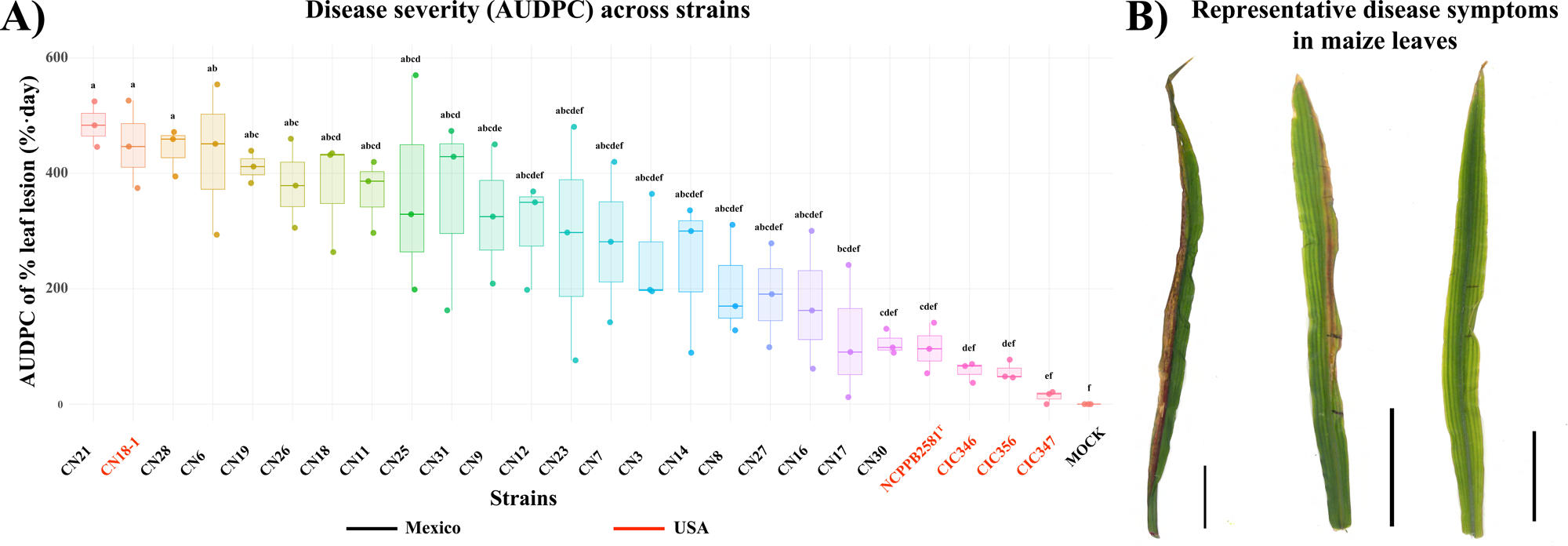
Virulence variation among *C. nebraskensis* strains in maize A632. A) Disease severity quantified as the area under the disease progress curve (AUDPC), calculated from the percentage of leaf lesion over time for each strain. Different letters denote statistically significant differences among strains (Kruskal–Wallis test followed by Dunn’s post hoc test, p < 0.05). Strain names are color-coded by country of origin. B) Representative disease symptoms observed on maize leaves following inoculation spanning the virulence spectrum. Black vertical bars indicate a scale of 5 cm.

### *C. nebraskensis* strain-specific growth dynamics in maize apoplastic fluid

To determine whether disease severity associates with apoplastic colonization capacity, *C. nebraskensis* growth was monitored in maize apoplastic fluid (Figure 4). Most strains exhibited biphasic growth characterized by an initial exponential phase, a plateau, and a second exponential phase, consistent with a diauxic-like transition (Figure 4A).

**Figure 4.**
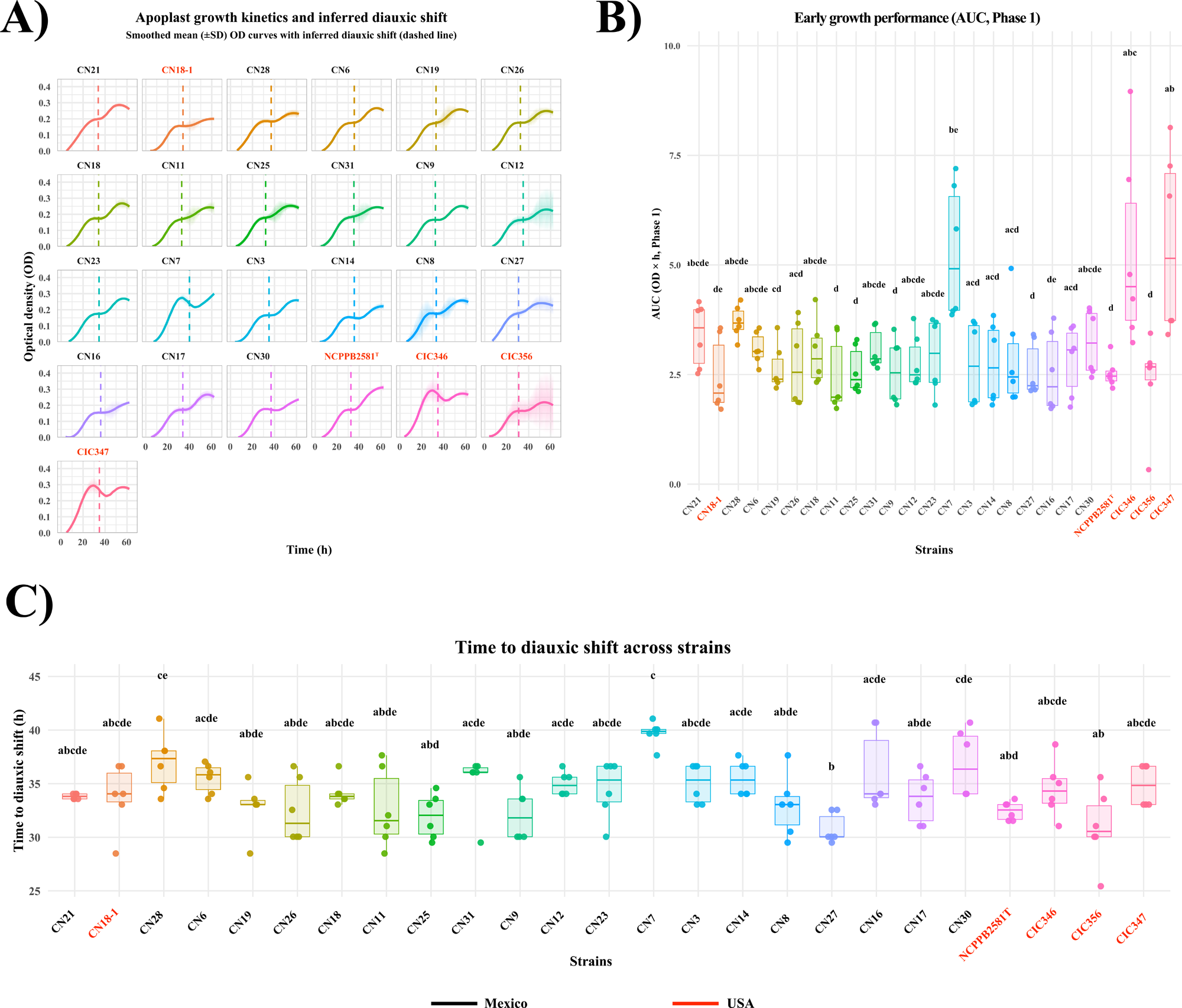
Apoplastic growth dynamics across strains. A) Apoplastic growth kinetics and inferred diauxic shift. Smoothed mean (±SD) optical density (OD) curves of strains grown in maize apoplastic fluid. Dashed vertical lines indicate the inferred diauxic shift, estimated from changes in growth slope over time. B) Early apoplastic growth performance. Area under the curve (AUDPC) during Phase I (pre-diauxic growth) calculated from apoplastic growth curves, summarizing early growth efficiency across strains. C) Distribution of time points at which the diauxic shift occurs during apoplastic growth.

Doubling time did not differ across strains in either phase (Phase 1: Kruskal–Wallis, p = 0.563; Phase 2: p = 0.414). In contrast, area under the growth curve (AUC) (Kruskal–Wallis, p = 1.68 × 10⁻⁴), and carrying capacity (K) (p = 0.00317) significantly varied among strains for phase 1 (Figure 4B; Supplementary Figure S2). Notably, the timing of the diauxic shift (t_break_) also varied across strains (Kruskal–Wallis, p = 1.27 × 10⁻⁵; Figure 4C), with some strains transitioning rapidly to phase 2, while others had prolonged exponential growth in phase 1, suggesting distinct patterns of apoplastic resource utilization (Figure 4).

### Enzymatic activity and genetic variation in carbohydrate- and protein-degrading traits across *C. nebraskensis* strains

To assess whether extracellular enzymatic functions could explain the differences in disease severity, we compared cellulase, proteases and amylase plate-based assays with genes sequence variation and genome-wide distributions. Cellulase and protease activities were different among strains (cellulase: χ² = 65.863, df = 24, p = 9.09 × 10⁻⁶; protease: χ² = 58.733, df = 24, p = 9.62 × 10⁻⁵), while no detectable starch degradation halos were observed for any strain, despite the presence of annotated amylase-related genes in the genomes (figure 5).

**Figure 5.**
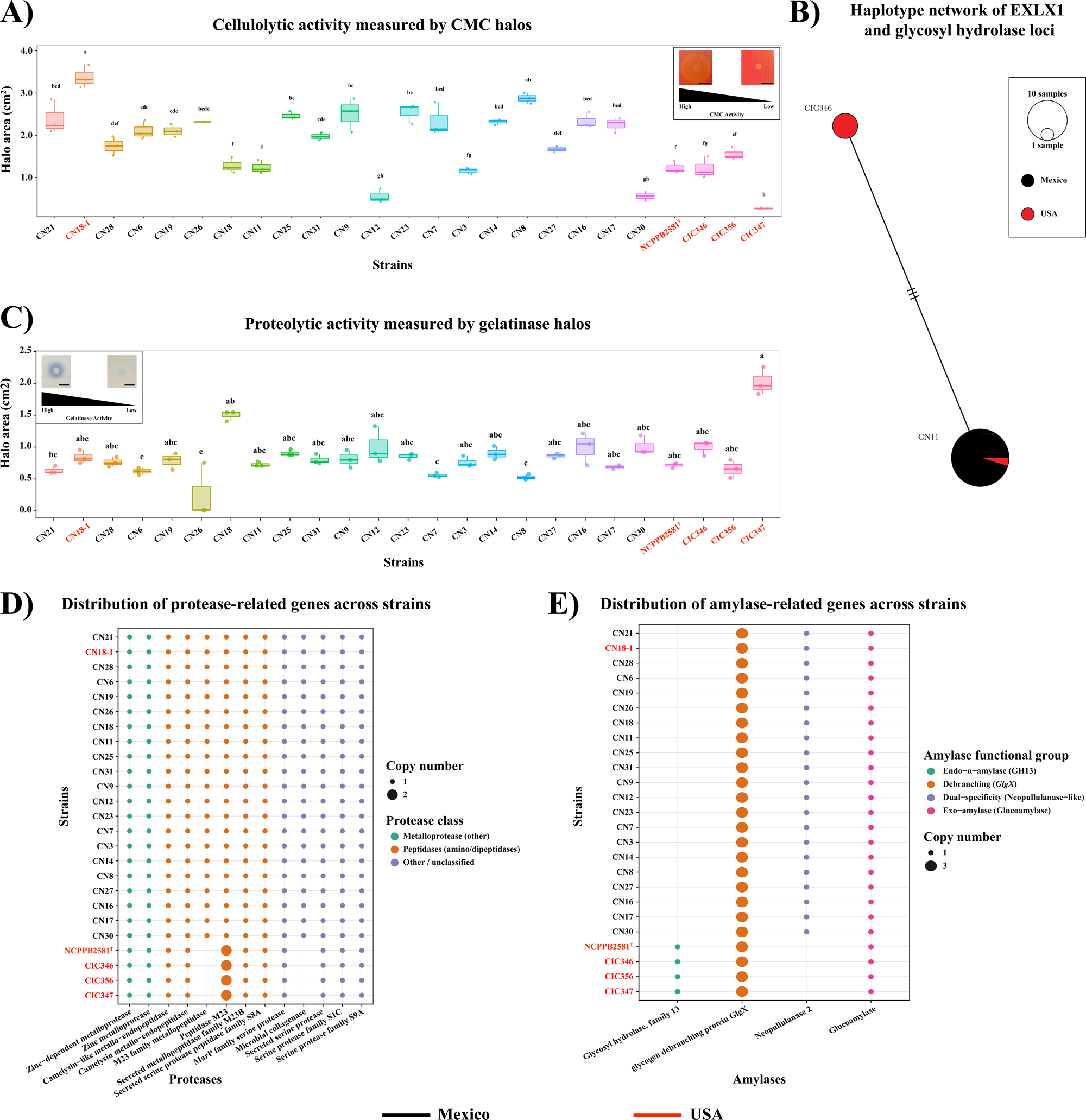
Extracellular enzymatic activity and associated functional gene content across strains. A) Cellulase activity quantified by halo area (cm.) measured on CMC agar plates. B) Cellulase activity haplotype network based on nucleotide variation in EXLX (expansin-like) and glycosyl hydrolase genes associated with cellulose degradation. Node size reflects the number of strains per haplotype, and colors indicate the country of origin. C) Protease activity quantified by gelatin hydrolysis through halo area (cm.) measured on gelatin agar plates, summarizing extracellular protease activity across strains. Statistical groupings are based on Dunn’s test with BH/FDR correction. D) Presence and copy number of annotated protease genes by functional class across strains. E) Presence and copy number of amylase and glycosyl hydrolase genes involved in starch and polysaccharide degradation across strains.

To determine whether variation in cellulase activity reflected gene SNP divergence, a haplotype network was constructed using two conserved loci, *exlX1* and a cellulase-family glycosyltransferase/cellulase-associated gene (Figure 5B). Nucleotide diversity across the concatenated alignment was low (π = 0.00043573), with three segregating and parsimony-informative sites detected. Tajima’s D was positive but not significant (D = 0.642, p = 0.2659), consistent with limited nucleotide sequence variation and no substantial departure from neutrality. Despite the low diversity, analysis of molecular variance revealed complete genetic partitioning among the predefined strain groupings (AMOVA: ΦST = 1.0, p < 0.001), indicating that the observed haplotypes were structured among populations.

Notably, four strains: CIC356, CIC347, CIC346, and the type strain NCPPB2581 shared a 462-nucleotide deletion spanning nucleotide positions 679 to 1140 within the *exlX1* gene. This deletion was absent from all other strains and resulted in a truncated *exlX1* allele. Strains carrying this deletion consistently associated with low to negligible CMC hydrolysis, providing a qualitative correspondence between structural variation at *exlX1* and cellulolytic activity. For protease-related genes, genome-wide annotation revealed heterogeneity for strains CIC356, CIC347, CIC346, and NCPPB2581 in the distribution of three genes (Figure 5E).

### Social behavior traits, bacteriocin activity, and extracellular matrix production

To explore whether virulence heterogeneity among *C. nebraskensis* strains could be attributed to social traits, we analyzed bacteriocin activity, extracellular polysaccharide (EPS) production, biofilm formation, and gene diversity associated with social behavior functions.

### Bacteriocin activity and associated genes diversity

Bacteriocin activity differed among strains and was influenced by growth medium (Figure 6A; File S3). Across all producer-indicator combinations, bacteriocin halos were wider on MBAL medium than on NBY (Kruskal–Wallis tests, p < 0.05; File S3). This indicates that bacteriocin expression and/or diffusion is enhanced under MBAL conditions, independent of strain identity. Among indicator strains, *Curtobacterium flaccumfaciens* pv. *flaccumfaciens* (VRP0036) exhibited consistent bacteriocin activity in both media, with mean inhibition halo areas in the intermediate-to-high activity group across multiple indicator strains (File S3). Notably, no strain exhibited broad-spectrum dominance across all indicators, underscoring strong specificity in producer-indicator interactions. The predicted bacteriocin biosynthetic gene clusters (BGCs) at the amino acid sequence level were conserved across strains, while the nucleotide level phylogenetic analysis revealed subgroup differentiation (Figure 6B). Mexican strains formed a clade with the U.S. strain from 1969, and the remaining U.S. strains clustered with the type strain NCPPB2581.

**Figure 6.**
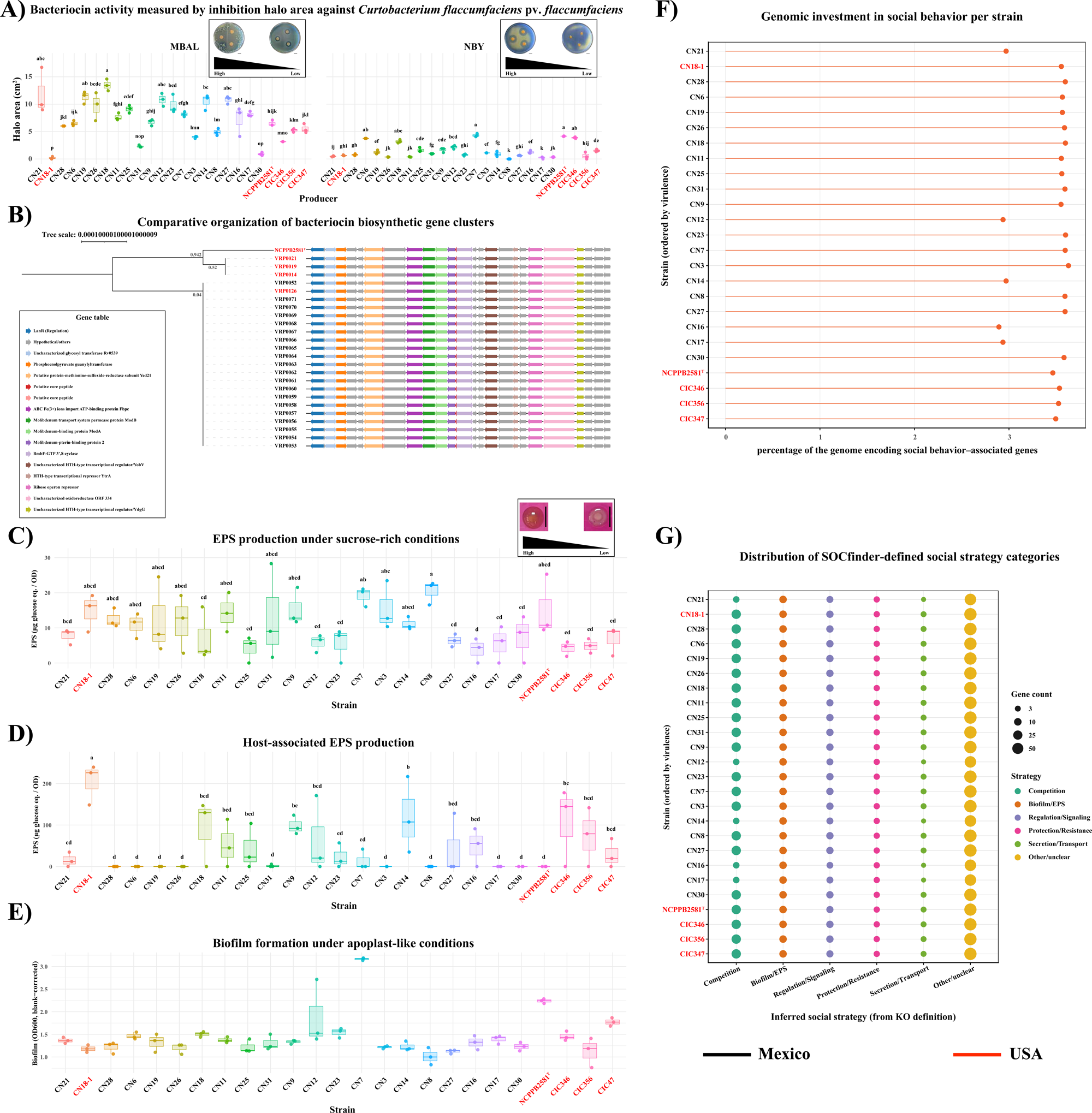
Social behavior and cooperative trait investment across *C. nebraskensis strains*. A) Bacteriocin activity quantified as inhibition halo area (cm.) against *Curtobacterium flaccumfaciens* pv. *flaccumfaciens* FH54, measured on MBAL and NBY media. Representative plates illustrate high- and low-inhibitory phenotypes; blue coloration results from a diffusible pigment produced by FH54 and is independent of bacteriocin activity. B) Phylogenetic reconstruction and gene content of bacteriocin Biosynthetic Gene Cluster (BGC) across strains, highlighting conservation and variation in regulatory, transport, and toxin-related genes. C) Exopolysaccharide (EPS) production in sucrose-amended medium, normalized by optical density (OD), showing strain-specific investment in extracellular matrix production. D) EPS production in maize apoplast extract, normalized by OD, reflecting hostrelevant extracellular matrix production under plant-mimicking conditions. E) Biofilm formation in maize apoplast extract quantified by crystal violet staining at OD₆₀₀, representing surface-associated growth under host-like conditions. F) Proportion of the genome composed of genes associated with social functions, including competition, biofilm/EPS production, regulation/signaling, protection/resistance, secretion/transport, and other cooperative traits. G) Proportion of the genome composed of genes associated with social functions, including competition, biofilm/EPS production, regulation/signaling, protection/resistance, secretion/transport, and other cooperative traits.

### Extracellular polysaccharide and biofilm

EPS production varied among strains and was dependent on growth conditions (Figure 6C–D). EPS production relative to biomass differed significantly among strains in sucrose-rich medium (Kruskal–Wallis: χ² = 42.006, df = 24, p = 0.01288) as well as in apoplast fluids (Kruskal–Wallis: χ² = 43.277, df = 24, p = 0.009246). Duncan’s multiple range test identified CN18-1 as the highest EPS producer under both conditions, followed by CN14, CIC346, CN9, and CN18. Notably, several strains that exhibited high EPS production under sucrose-rich conditions (e.g., CN7 and CN8) produced moderate to low levels of EPS in apoplast fluids, indicating condition-dependent regulation of EPS synthesis.

Biofilm formation also varied significantly among strains (Figure 6E; Kruskal–Wallis: χ² = 55.9, df = 24, p = 0.000234). Strains exhibiting elevated EPS production under sucrose-rich conditions frequently showed increased biofilm formation, whereas EPS production in apoplast fluids did not display a consistent correspondence with biofilm levels. Together, these results suggest that EPS produced under carbohydrate-rich conditions may contribute more directly to biofilm formation than EPS produced under host-associated conditions.

### Genomic basis of social behavior heterogeneity

Genome-wide profiling revealed heterogeneity across strains in both the relative genomic investment and the functional composition of social behavior traits (Figure 6). Strains distributed along a broad gradient in the proportion of genes annotated to social behavior categories, ranging from low to high allocation (Figure 6F). No single functional class accounted for this variation; instead, heterogeneity reflected cumulative differences across multiple categories, including bacteriocins, extracellular polysaccharide-related functions, secretion-associated genes, and other interaction-related traits. Notably, genomic allocation did not consistently predict phenotypic output (Figure 6G). Several strains with elevated social behavior gene content showed modest phenotypic expression, while others displayed strong social phenotypes despite limited genomic investment.

### Pigment-associated phenotypes and genetic variation in environmental protection traits

Bacterial pigments function in protection against environmental stressors (oxidative damage, antimicrobial activity, UV radiation, nutrient limitation) and may also modulate social interactions through phenotypic signaling. Thus, pigmentation variation reflects both ecological resilience and social trait expression. We characterized strain-level pigmentation variation through absorbance spectral profiles and genomic analysis of terpene and thiamine biosynthesis (Figure 7).

**Figure 7.** Pigment production and thiamine-associated genetic traits linked to environmental protection. A) Hierarchical clustering of pigment phenotypes. Dendrogram constructed using Euclidean distances derived from quantitative absorbance measurements of pigment production. The inner ring represents colony color categories extracted from digital images of colonies grown on NBY agar. The outer ring displays representative colony images for each strain grown on NBY agar. B) Haplotype network based on nucleotide variation in terpene precursor (region 6) and terpene biosynthetic (region 9) loci identified by antiSMASH. Node size reflects the number of strains per haplotype, and colors correspond to pigment phenotype classes defined in panel A. C) Presence and copy number of genes involved in thiamine biosynthesis, transport, and regulatory functions across genomes.

### Spectral characterization and genomic comparisons

Spectral absorbance profiles (200–1,000 nm) displayed strain-specific variation, with multiple peaks primarily detected in the UV and near-visible ranges (Figure 7A). Clustering strains based on Euclidean distances among full spectral curves identify four pigment-absorbance categories: high, intermediate, moderate, and low, indicating that pigmentation differences are reflected by the spectral profile. To evaluate whether pigment-associated groupings aligned with genetic variation in terpene biosynthesis, we constructed a haplotype network using nucleotide sequences from the terpene precursor (region 6) and terpene biosynthetic (region 9) regions identified by antiSMASH (Figure 7B). The concatenated alignment showed nucleotide diversity (π = 0.00245533), with 537 segregating sites and 525 parsimony-informative sites. Tajima’s D was positive but not significant (D = 0.1317, p = 0.4376), consistent with no strong deviation from neutral expectations at these loci. Analysis of molecular variance indicated no significant population structure (ΦST = −0.0201, permutation p ≈ 0.44), suggesting that terpene-region sequence variation is broadly shared across strains. We analyzed distribution of thiamine biosynthesis genes as thiamine is essential for orange pigment production in *C. nebraskensis* (Bradbury 1991). Genome-wide annotation revealed heterogeneity in the distribution and copy number of genes related to thiamine metabolism and transport (Figure 7C). While several thiamine-associated genes were conserved across all strains, others had variations in specific strain subsets. Notably, these differences partially overlapped with the pigment-intensity categories defined by spectral clustering (Figure 7A, C).

## Discussion

This study demonstrates that *Clavibacter nebraskensis* is an ecologically heterogeneous species structured around alternative survival and competition strategies. By integrating phylogenomics and pangenomic analysis, host-interaction assays, enzymatic activity profiling, and social behavior characterization across strains, we provide the most comprehensive ecological and genomic framework to date for *C. nebraskensis*.

Whole genome phylogenomics confirmed that *C. nebraskensis* forms a monophyletic species clade that segregates by geographic origin. Pangenome reconstruction revealed that this segregation is not driven by core genome but rather by variations in accessory gene content, a pattern reported in other microbial systems (Bertazzoni et al. 2018; Chang and McGwire 2002; Croll and McDonald 2012). Specifically, Mexican strains and the 1969 U.S. strain share a distinctive accessory genetic signature that diverges from other *C. nebraskensis* isolates. This pattern reflects gene loss and recombination rather than ongoing adaptive acquisition, suggesting that *C. nebraskensis* is a closed genome and that strain diversification operates through modulation of existing accessory genes (Rosconi et al. 2022; Rouli et al. 2015). Notably, the structured genetic diversity observed in Mexican populations, indicates endemic populations rather than recent introductions, consistent with the genomic structure of widely distributed plant pathogens (Flores-López et al. 2024; Morales-Galván et al. 2022; Robinson et al. 2011; Sarkar and Guttman 2004; Smith et al. 2000; Timilsina et al. 2022).

Symptom severity and lesion expansion among *C. nebraskensis* strains ranged from extensive necrosis with high AUC values to mild water soaking or freckling. Consistent with previous reports, virulence variations in *C. nebraskensis* were independent of genetic diversification (Agarkova et al. 2011; Webster et al. 2020). We next assessed whether virulence could be explained by differences in maize apoplastic growth capacity. Although all strains exhibited biphasic growth, virulence was not determined by maximum growth rates. Instead, virulence appeared linked to early biomass accumulation and the timing of the diauxic transition, metrics reflecting early nutrient-exploitation strategies and metabolic flexibility that better explain disease progression (Bhagwat et al. 2025; Khokhani et al. 2017). We acknowledge that virulence assessments were conducted using the maize inbred line A632. Testing additional U.S. and Mexican maize varieties could reveal genotype-dependent virulence variation not captured in this study.

From enzymatic assays, only cellulases and extracellular proteases varied among *C. nebraskensis* strains; amylase activity was not detected in any strain. Cellulase activity correlated positively with virulence for some strains, consistent with studies showing that endoglucanase deletion reduces symptoms (Lin et al. 2021b; Veregge et al. 2025). In contrast, extracellular protease activity showed no clear correlation with virulence. When looking at the genotype–phenotype relationships, cellulase-associated genes showed minimal sequence divergence across strains clustering primarily by geographic region. In contrast, extracellular protease genes exhibited substantial variation in copy numbers and functional annotation. Notably, amylase-associated genes were conserved across all strains despite the absence of amylolytic activity in the assays. The absence of genotype–phenotype direct correlations demonstrates that *C. nebraskensis* pathogenesis could be regulated by factors such as transcriptional regulation, inactive under the assay conditions, protein secretion efficiency, environmental responsiveness, and substrate recognition (Lee Erickson and Schuster 2024; Lin et al. 2021b; Roussin-Léveillée et al. 2024; Veregge et al. 2025; Zimmerman et al. 2013).

Social behavior traits described here as bacteriocins, EPS and biofilm production, further demonstrated that virulence in *C. nebraskensis* is not a simple function of cooperative or communal investment. Bacteriocin activity was widespread across *C. nebraskensis* strains, yet activity did not correlate with virulence and inhibiting activity was restricted to subsets of closely related strains, consistent with bacteriocins function as narrow, strain-specific suppressors (Gross and Vidaver 1979; Smidt and Vidaver 1986; Vidaver 1983, 2002; Vidaver and Buckner 1978). Moreover, bacteriocin production and diffusion were strongly influenced by the growth medium, as reported before by Gross and Vidaver (1979). MBAL medium, which approximates plant-derived substrates more closely than NBY, showed more discriminatory bacteriocin activity across strains, suggesting that bacteriocin expression in MBAL may better reflect competitive dynamics in planta (Ehau-Taumaunu and Hockett 2022). However, direct assessment of bacteriocin activity in apoplast-mimicking media will be required to validate this hypothesis. Finally, nucleotide-level analysis of bacteriocin BGC revealed clear subgroup differentiation. This pattern suggests that evolutionary divergence acts primarily on regulatory control rather than on the protein cluster. Such regulatory divergence is consistent with bacteriocins fine-tuning competitive interactions within strain communities rather than determining virulence outcomes. (Acedo et al. 2018; Backman et al. 2024; Reuben and Torres 2024; Rooney et al. 2020; Rutter et al. 2024; Sharp and Foster 2025).

Extracellular polysaccharide (EPS) production and biofilm formation were also influenced by the medium. EPS levels in each strain changed between apoplast and sucrose-rich environment, while for biofilm formation some strains were consistent in both media. Biofilm as well as EPS production did not show a correlation with pathogenicity, like other plant pathogenic bacteria (Gajdács et al. 2021; Heredia-Ponce et al. 2020; O’Banion et al. 2025; Tkaczyk 2025). At the genomic analysis, strains classified within higher social-investment categories did not consistently overlap with those exhibiting the strongest bacteriocin production, EPS accumulation, or biofilm formation, underscoring that social behavior is distributed across partially independent multigenetic modules rather than coordinated as a single trait.

Pigment-associated traits provided additional evidence that diversification in *C. nebraskensis* is not associated with virulence. Strains exhibited distinct spectral absorbance profiles clustering into discrete pigment-intensity categories independent of phylogeny or disease severity. We also observed that pigmentation in *C. nebraskensis* is dependent on thiamine availability (Bradbury 1991). Colonies were consistently white to cream on thiamine-free PDA, whereas thiamine supplementation promoted growth and induced the characteristic orange pigmentation. Genomic analysis revealed that terpene biosynthetic genes showed moderate nucleotide diversity, while thiamine biosynthesis and transport genes exhibited presence/absence variation and copy number differences, yet these genetic variations did not consistently predict pigmentation phenotypes. In plant-associated bacteria broadly, pigmentation-associated traits have been linked to tolerance to environmental stressors, such as oxidative stress, UV radiation, and nutrient limitation, promoting survival on plant surfaces and under unfavorable conditions, rather than directly serving as determinants of virulence (Armstrong 1994; Costliow and Degnan 2017; Huang et al. 2024; Ram et al. 2020; Rodríguez-Villalón et al. 2008). (Costliow and Degnan 2017; He et al. 2020; Jacobs et al. 2005). In other bacterial phytopathogens, pigmentation directly contributes to pathogenic fitness (Mohammadi et al. 2012; Rodríguez-Mejía 2010). The diversity of pigmentation functions across bacterial species indicates that these associations are pathogen-specific, determined by unique combinations of host genetics, pathogen genetics, and environmental context rather than representing a universal virulence mechanism.

Based on our findings, we propose two ecological strategies for *C. nebraskensis* (Figure 8). Strategy I (“niche engineers”) prioritizes long-term survival through asymptomatic colonization of crop residues and epiphytic surfaces, enabling overwintering and maintenance of populations. This strategy aligns with epidemiological evidence showing that *C. nebraskensis* is rarely seed-transmitted and that outbreaks result from seasonal buildup of inoculum until reaching a critical threshold for symptom development (Block et al. 2019). Strategy II (“aggressive competitors”) operates once sufficient inoculum is present, disrupting this balance through rapid exploitation of host tissue and competitive exclusion within the plant microbiome. Rather than operating solely as direct virulence factors, the observed strain-specific variation in cellulase, protease, and bacteriocin activity functions as a dual mechanism: causing direct host tissue damage while simultaneously changing the microbiota and promoting dysbiosis (Backman et al. 2024; DeBoy et al. 2008; Ehau-Taumaunu and Hockett 2022; Leveau 2024; Peyraud et al. 2016; Pfeilmeier et al. 2024; Pinto et al. 2025).

**Figure 8.**
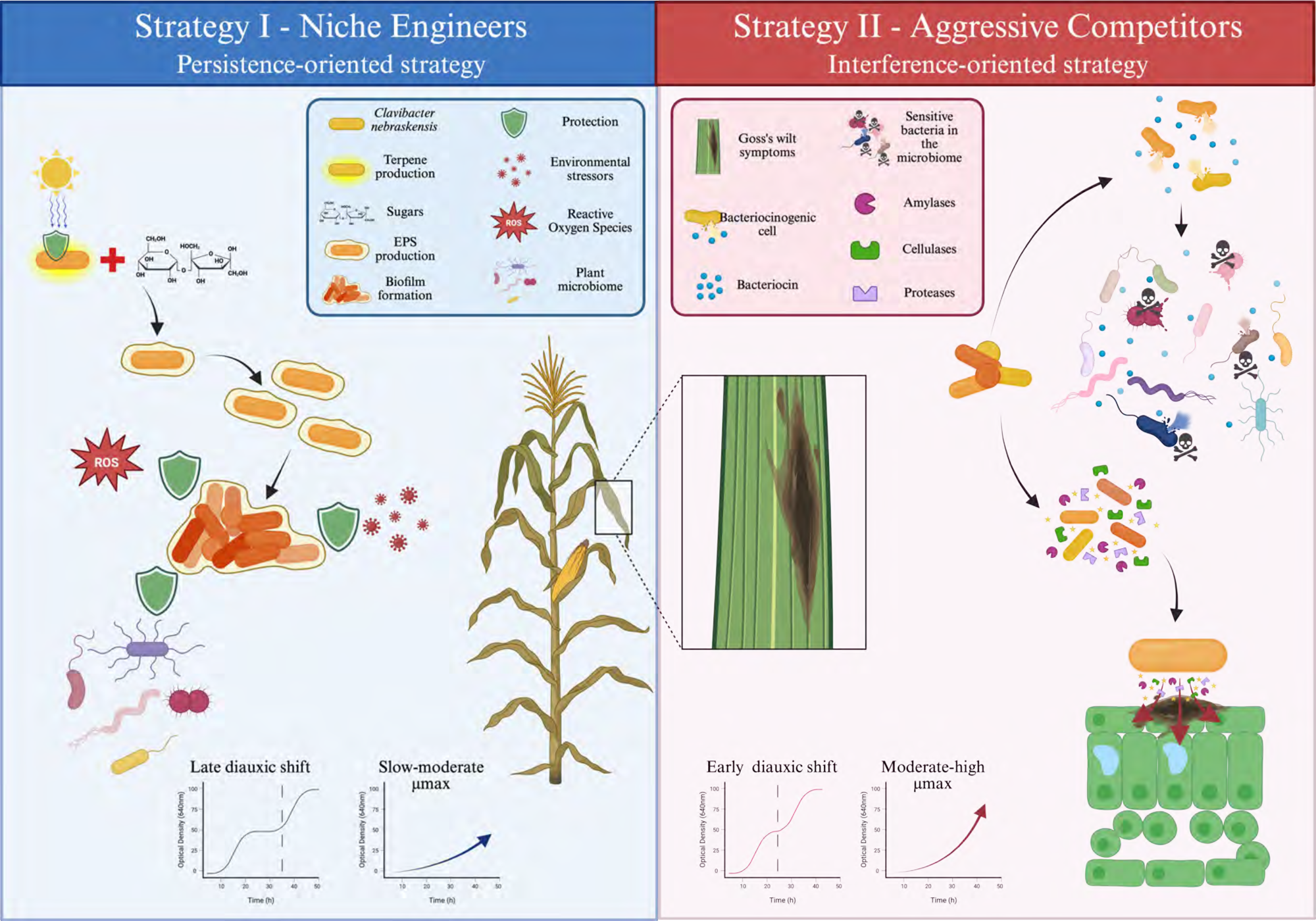
Conceptual model of alternative fitness strategies in *Clavibacter nebraskensis*. Schematic summary of two dominant fitness strategies inferred from phenotypic assays, genomic analyses, and apoplastic growth experiments. Strategy I – Niche engineers. Strains following this strategy emphasize persistence within the maize apoplast through production of protective traits, including exopolysaccharides (EPS), biofilm formation, and terpene-associated pigments. These traits contribute to tolerance of environmental stressors such as reactive oxygen species and interactions with the resident plant microbiome. Apoplastic growth is characterized by delayed diauxic shifts and slow-to-moderate maximum growth rates (μmax), consistent with sustained resource utilization and niche stabilization. Strategy II – Aggressive competitors. Strains following this strategy prioritize rapid exploitation and competitive exclusion through bacteriocin production and secretion of extracellular enzymes, including cellulases, amylases, and proteases. These traits contribute to microbial antagonism and host tissue degradation. Apoplastic growth is characterized by earlier diauxic shifts and moderate-to-high μmax, consistent with rapid resource use and increased symptom development.

Strategies I and II are complementary ecological approaches representing alternative allocations of metabolic and regulatory resources with trade-offs between persistence and competitive aggressiveness. These strategies could help explain Goss’s wilt dynamics: when conditions favor Strategy I, *C. nebraskensis* populations remain cryptic and undetectable through traditional surveillance. When environmental or agronomic conditions trigger Strategy II, resident populations transition to pathogenic expression. Consequently, the absence of detected disease in a particular year or survey area does not indicate pathogen absence but rather reflects ecological conditions suppressing outbreaks (Jackson et al. 2007). Furthermore, these strategies could explain why seed transmission is low (Block et al. 2019). Finally, the presence of *C. nebraskensis* in Tlaxcala could be consistent with Strategy I, where the bacteria have persisted through asymptomatic colonization, ruling out recent introduction. (Flores-López et al. 2024; Janzen et al. 2022; Li 2024; Louette and Smale 2000; Sawers 2011).

Our results demonstrate that effective pest risk assessment and disease management for bacterial pathogens require integration of phylogenomics, population ecology, and agroecosystem context. Single-year surveillance fails to detect cryptic persistence, rare transmission events, and environmentally contingent disease expression, leading to underestimated risk and mischaracterized epidemiology. Evaluation of introduced maize genotypes and new varieties needs to be monitored. Management strategies should adopt multi-year, ecologically informed approaches that account for these dynamics to accurately assess risk and implement sustainable control for *C. nebraskensis* and emerging bacterial pathogens.

## Acknowledgments

The authors gratefully acknowledge Dr. Elizabeth Rogers and Aaron Sechler (USDA-ARS, Foreign Disease/Weed Science Research Unit) for providing bacterial strains used as indicators in the bacteriocin assays, including the type strain *Clavibacter nebraskensis* FH36 (NCPPB2581). We thank Dr. Tamra Jackson-Ziems (University of Nebraska–Lincoln) for providing strain CN18-1 (1969) from the original collection of Dr. Anne Vidaver.

## Data availability

All genome sequences were submitted to NCBI under Bioproject PRJNA1266758.

