## Supplementary material for "Ecogenomic Diversity of *Clavibacter nebraskensis* in North America": File

Table S1

| | | | |
|---|---|---|---|
| ATCC33566 | *Clavibacter tessellarius* | 1978 USA | GCF_002240635.1 |
| DM1 | *Clavibacter zhangzhiyongii* | 2017 Australia | GCF_014775655.1 |
| PF008 | *Clavibacter capsici* | 1999 South Korea | |
| CFBP8615 | *Clavibacter lycopersici* | 2015 Iran | |
| CFBP8217 | *Clavibacter phaseoli* | 2007 Chile | GCF_021923065.1 |
| HF4 | *Clavibacter nebraskensis* | 2012 USA: Iowa | GCF_009739595.2 |
| SL1 | *Clavibacter nebraskensis* | 1983 USA: Iowa | GCF_023277825.1 |
| NCPPB2581T | *Clavibacter nebraskensis* | 1971 USA: Nerbaska | GCF_000355695.1 |
| DOAB395 | *Clavibacter nebraskensis* | 2014 Canada: Manitoba | GCF_001643055.1 |
| DOAB397 | *Clavibacter nebraskensis* | 2014 Canada: Manitoba | GCF_000966585.1 |
| 06-1 | *Clavibacter nebraskensis* | 2006 USA: Iowa | GCF_009739635.2 |
| 7580 | *Clavibacter nebraskensis* | 2006 USA: Iowa | GCF_009739615.2 |
| CNK2 | *Clavibacter nebraskensis* | 1972 USA: Kansas | GCF_023539135.1 |
| CIBA | *Clavibacter nebraskensis* | 1996 USA: Nerbaska | GCF_023277845.1 |
| STE U9949 | *Clavibacter nebraskensis* | 2024 South Africa | GCF_051046835.1 |
| STE U9951 | *Clavibacter nebraskensis* | 2024 South Africa | GCF_051046825.1 |
| STE U9948 | *Clavibacter nebraskensis* | 2024 South Africa | GCF_051046845.1 |
| A6096 | *Clavibacter nebraskensis* | 1971 USA: Nerbaska | GCF_021923125.1 |
| 419B | *Clavibacter nebraskensis* | 2011 USA: Nerbaska | GCF_023277865.1 |
| CN205 | *Clavibacter nebraskensis* | 2024 South Africa | CP173672.1 |
| 44 | *Clavibacter nebraskensis* | 2011 USA: Nerbaska | GCF_023279165.1 |
| ATCC10253 | *Clavibacter insidiosus* | 1960 USA | GCF_003076355.1 |
| CFBP2404 | *Clavibacter insidiosus* | 1955 USA | GCF_003693415.1 |
| LMG3663 | *Clavibacter insidiosus* | 1955 USA | GCF_002240565.1 |
| ATCC33113 | *Clavibacter sepedonicus* | 1968 Canada | GCF_000069225.1 |
| LMG7333 | *Clavibacter michiganensis* | 1957 Hungary | GCF_021216655.1 |
| VKMAc1403 | *Clavibacter michiganensis* | 1957 Hungary | GCF_900168345.1 |
| A6099 | *Clavibacter seminis* | 2013 USA | GCF_021919125.1 |
| CFBP8216 | *Clavibacter californiensis* | 2000 USA | GCF_021952865.1 |
| CP139624 | *Clavibacter quasicaliforniensis* | 1998 USA | GCF_051201715.1 |

Table S2

| Strain | VRP ID | Species | Donnor |
|---|---|---|---|
| FH53 | VRP0035 | Curtobacterium flaccumfaciens pv. flaccumfaciens | Elizabeth Rogers, USDA |
| FH54 | VRP0036 | Curtobacterium flaccumfaciens pv. flaccumfaciens | Elizabeth Rogers, USDA |
| CO103 | VRP0040 | Curtobacterium flaccumfaciens pv. oortii | Elizabeth Rogers, USDA |
| FH34 | VRP0041 | Curtobacterium flaccumfaciens pv. oortii | Elizabeth Rogers, USDA |
| FH8 | VRP0042 | Curtobacterium flaccumfaciens pv. betae | Elizabeth Rogers, USDA |
| FH7 | VRP0051 | Clavibacter tessellarius | Elizabeth Rogers, USDA |

### A) Pangenome accumulation

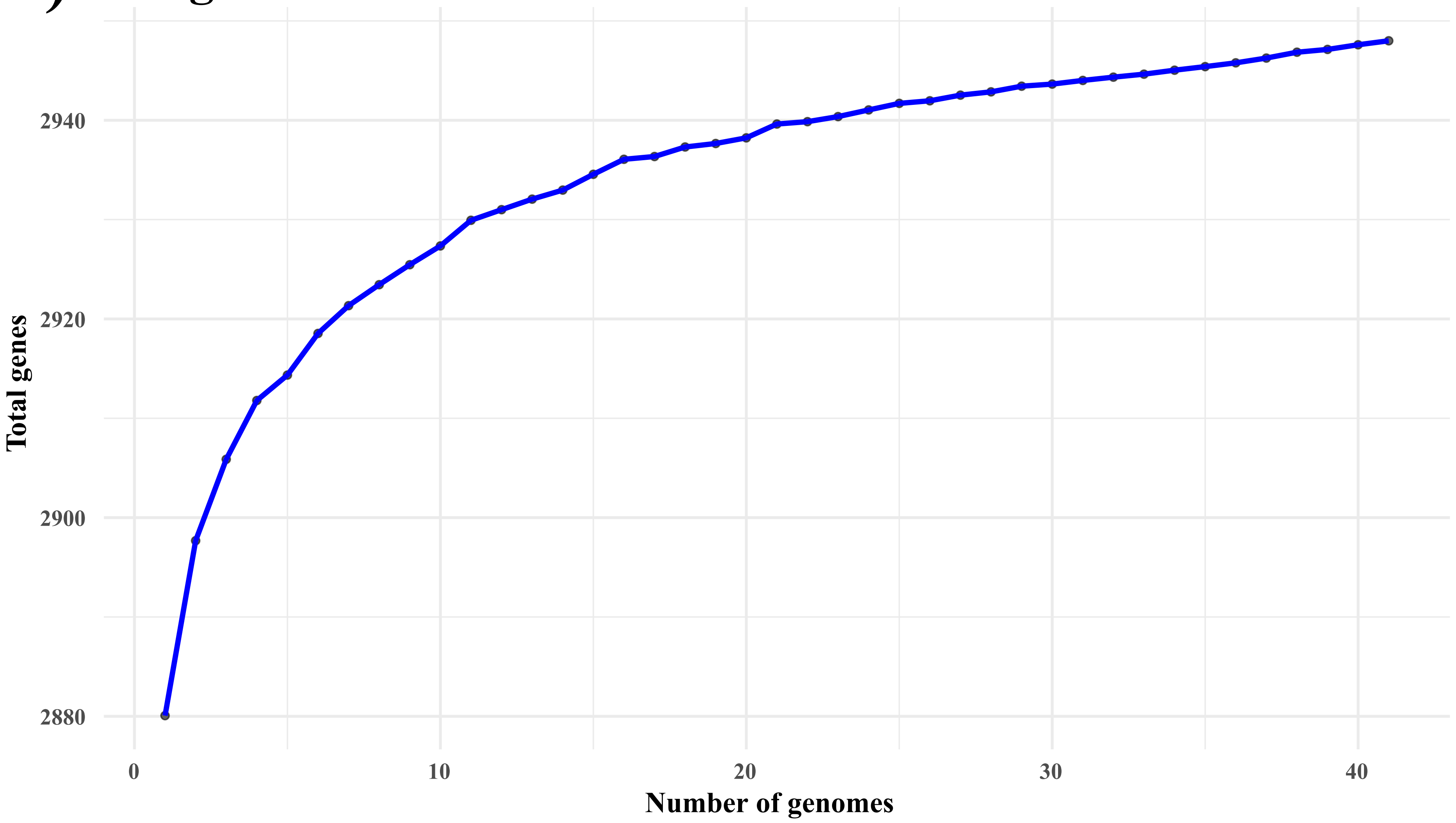


### B) New genes per genome

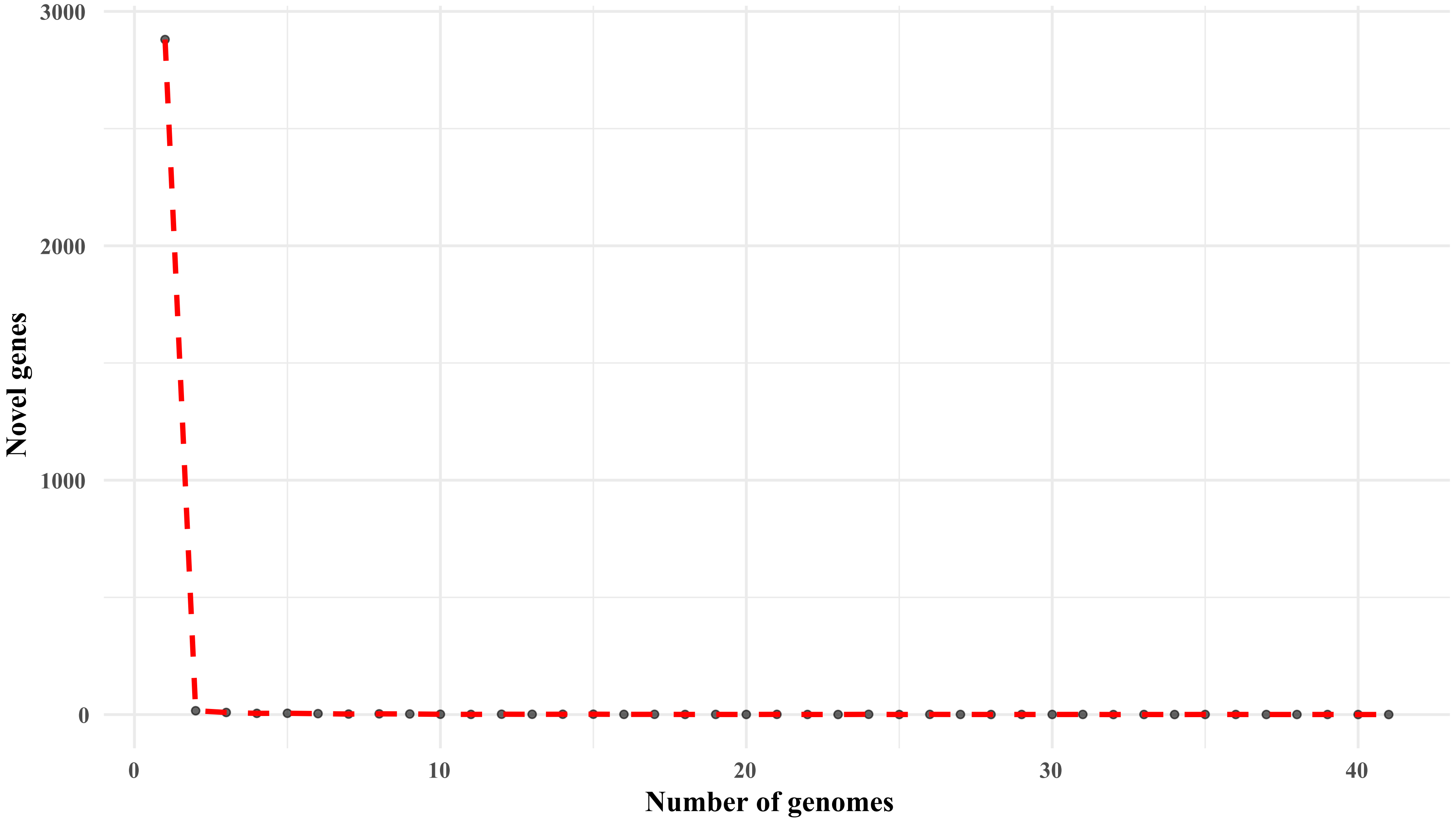

SupFile2

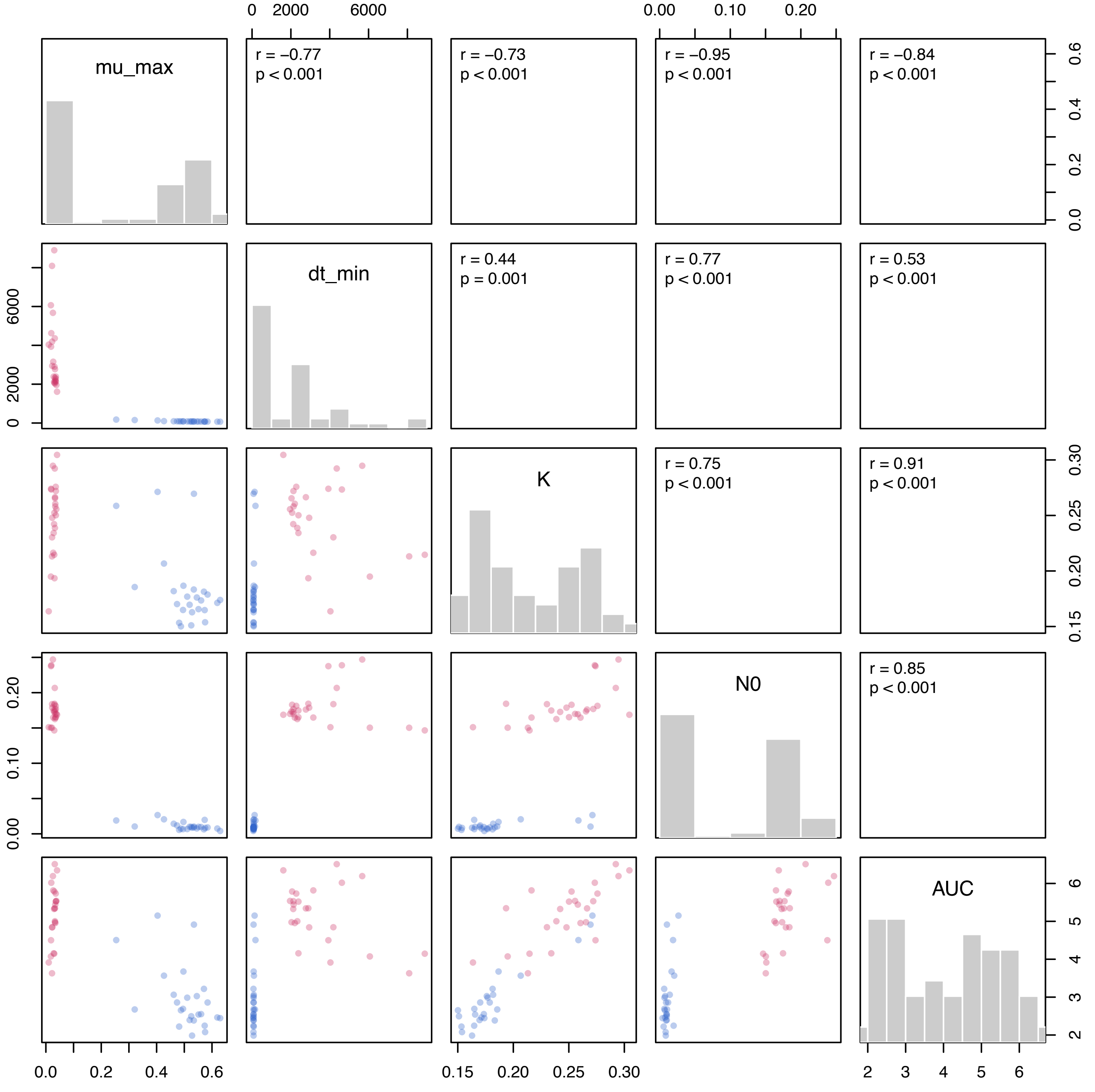

mu_max
dt_min
K
N0
AUC
r = −0.77
p < 0.001
r = −0.73
p < 0.001
r = −0.95
p < 0.001
r = −0.84
p < 0.001
r = 0.44
p = 0.001
r = 0.77
p < 0.001
r = 0.53
p < 0.001
r = 0.75
p < 0.001
r = 0.91
p < 0.001
r = 0.85
p < 0.001
0 2000 6000
0.00 0.10 0.20
0.0 0.2 0.4 0.6
0.15 0.20 0.25 0.30
2 3 4 5 6

supfile3

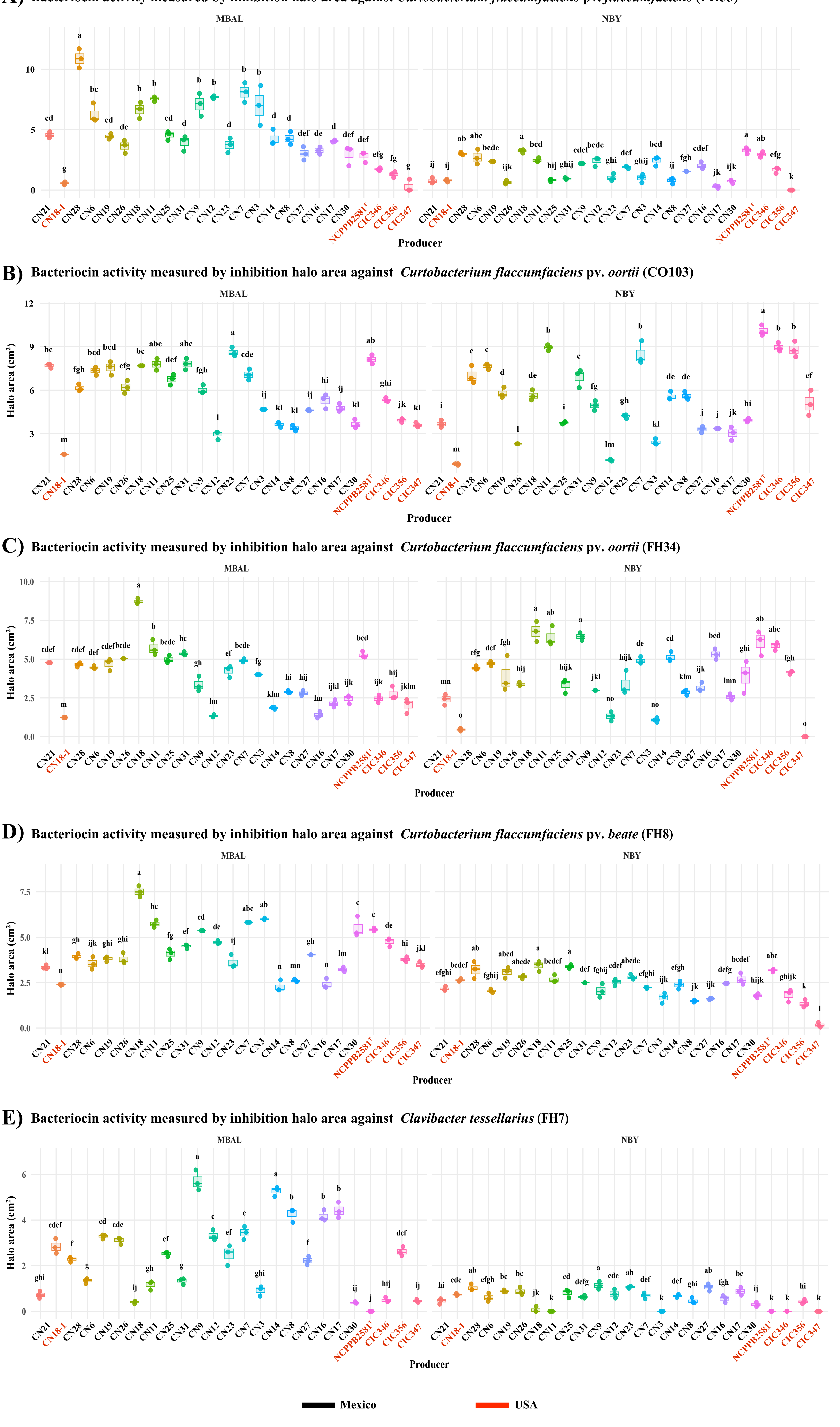
A) Bacteriocin activity measured by inhibition halo area against Curtobacterium flaccumfaciens pv. flaccumfaciens (FH53)
B) Bacteriocin activity measured by inhibition halo area against Curtobacterium flaccumfaciens pv. oortii (CO103)
C) Bacteriocin activity measured by inhibition halo area against Curtobacterium flaccumfaciens pv. oortii (FH34)
D) Bacteriocin activity measured by inhibition halo area against Curtobacterium flaccumfaciens pv. beate (FH8)
E) Bacteriocin activity measured by inhibition halo area against Clavibacter tessellarius (FH7)
MBAL
NBY
Halo area (cm²)
Producer
CN21
CN18-1
CN28
CN6
CN19
CN26
CN18
CN11
CN25
CN31
CN9
CN12
CN23
CN7
CN3
CN14
CN8
CN27
CN16
CN17
CN30
NCPPB2581ᵀ
CIC346
CIC356
CIC347
Mexico
USA
